# Mechanism of substrate binding by the SARS-CoV-2 NiRAN domain and modulation of its activities during replication

**DOI:** 10.1101/2025.10.31.685727

**Authors:** Ashwin Uday, Muhammed Navas, Pazhayamadathil Sathyapalan Sivaprasad, Ufaq Bhat, Kayarat Saikrishnan

## Abstract

The SARS-CoV-2 Nidovirus RdRp-associated nucleotidyltransferase (NiRAN) domain initiates viral genome capping by RNAylating nsp9 with the 5′-pppA-end of the genome followed by GDP-dependent deRNAylation to form the core capped GpppA-genome. Additionally, it cycles nsp9 through NMPylation-deNMPylation to generate GpppN. It is unclear how the distinct substrates, 5′-pppA-RNA and NTP, are bound, and how NiRAN balances RNAylation versus NMPylation. Earlier models proposed a common base-up pose for both the substrates. Here, structure-guided mutagenesis and reconstitution assays show that 5′-pppA indeed binds base-up during RNAylation, revealing that nsp12-Asp711 confers adenine selectivity, whereas, NTP adopts a perpendicular base-out pose during NMPylation. NiRAN intrinsically favors NMPylation over RNAylation, but nsp13 NTPase activity flips this preference. RNAylation weakens when RdRp is RNA-bound or replicating it, suggesting that a trans-acting NiRAN associated with an RNA-free RdRp performs capping. These findings provide insights into the orchestration of NiRAN activities and potential druggable sites for anti-viral therapeutics.

## Introduction

SARS-CoV-2, the etiological agent of COVID-19, triggered a global pandemic that tremendously impacted human health and socioeconomic systems^1,2^. SARS-CoV-2 belongs to the family *Coronaviridae* within the order *Nidovirales*^3^. As new variants of SARS-CoV-2 and other coronaviruses emerge, these continue to pose a serious threat to human life and society and there is an urgency to study them in detail^4–6^. SARS-CoV-2 possesses a large genome (∼30kb) of polycistronic single-stranded RNA^7^. This genome codes for 16 non-structural proteins (nsps) from the first open reading frame (ORF), followed by multiple structural as well as accessory proteins from the remaining ORFs^8^. Many of the nsps together form the machinery required for viral genome replication and mRNA (sub-genomic RNA) transcription^7^.

At the core of the replication machinery is nsp12, which contains the C-terminal RNA dependent RNA polymerase (RdRp) domain. nsp7 and nsp8 are essential accessory proteins that together with nsp12 form the core replication transcription complex (RTC) in the stoichiometric ratio 1:2:1, respectively^9,10^. RTC is the minimal machinery that can replicate RNA. N-terminus of nsp12 contains the nidovirus RdRp-associated nucleotidyltransferase (NiRAN) domain that is specialized to initiate capping of the viral RNA^11–13^. Capping of the replicated RNA with 5’-methylated guanosine is essential for its stability, protection from nucleases and translation initiation in a eukaryotic host^14,15^. Consequently, it is an important process for viral genome replication and the viral life cycle. Co-occurrence of polymerase with nucleotidyltransferase within a single protein is a unique hallmark and a molecular marker of the order *Nidovirales*^11,16^.

Previous studies have shown NiRAN is a multifunctional enzyme that catalyzes the first two steps of RNA capping^17^ (Fig. 1A). The first step is the transfer of 5’-monophosphate RNA (5’-pA-RNA) from 5’-triphosphate RNA, 5’-pppA-RNA, to the main chain amino group of the N-terminal asparagine of nsp9, forming a phosphoramidite bond and release of pyrophosphate in a reaction termed as RNAylation. The 5’-nucleotide of the RNA has to be an adenosine triphosphate, in the absence of which RNAylation does not occur^17^. In the second step, the 5’-pA-RNA covalently bonded to nsp9 is transferred to the β-phosphate of GDP bound to NiRAN. In this GDP-polyribonucleotidyltransferase reaction, referred to as deRNAylation, free nsp9 and GpppA-RNA are released, forming the core cap structure. The GpppA-RNA is subsequently modified by nsp14 and nsp16, which methylate the N7 of the guanine base and 2’O of the first adenosine nucleoside, respectively, leading to the formation of the complete cap structure – ^7Me^GpppA_2’-OMe_-RNA^18,19^. In addition to 5’-pppA-RNA, NiRAN can also use nucleoside triphosphate (NTP) to transfer nucleoside monophosphate (NMP) to nsp9 forming NMP-nsp9 (NMPylation)^20,21^. In presence of GDP, NMP gets transferred from nsp9 forming GpppN (deNMPylation)^21^. The physiological role of NMPylation is not known, but it is speculated that it could be involved in preventing nsp9 degradation and/or protein priming for RNA extension^11,22–24^.

**Figure 1:**
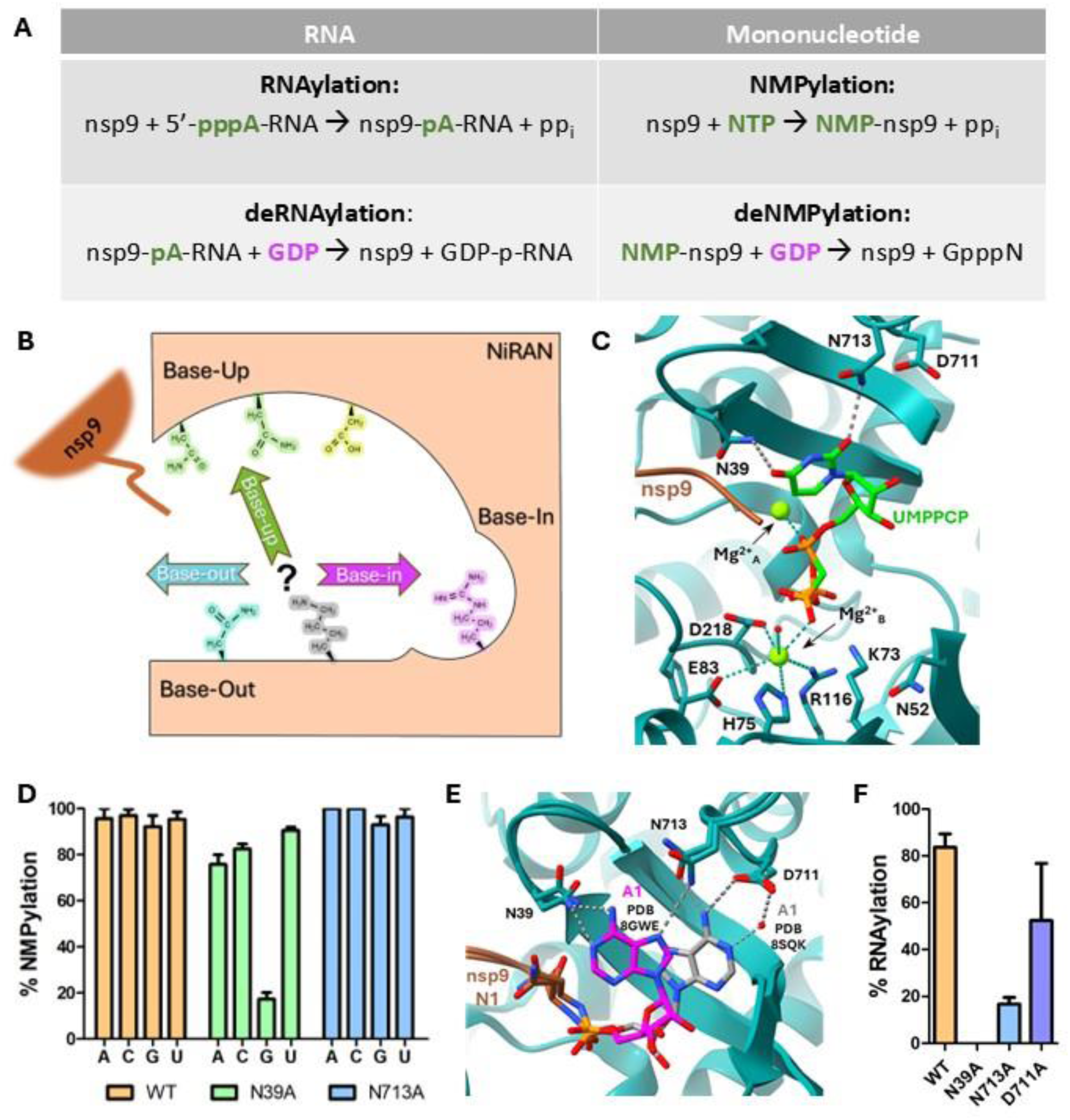
Base-up pose the nucleotide binding mode during RNAylation. (A) Reaction schema of the four reactions catalyzed by SARS-CoV-2 NiRAN domain. Nucleotides are colored as per their poses during the reaction attributed prior to this study. Base-up in green and base-in in magenta. (B) Schematic showing nucleotide binding poses in NiRAN. (C) Cartoon representation of UMPPCP bound to the NiRAN active site in the base-up pose (PDB ID 8SQ9). Some of the interactions made by NiRAN residues with the nucleotide and Mg^2+^ are shown. (D) NMPylation activity of wild type (WT) RTC and RTC with indicated mutants of nsp12 using the nucleotides ATP (A), GTP (G), CTP (C) and UTP (U). The reaction products were analyzed using SDS-PAGE and Coomassie staining and quantified using gel densitometry. %NMPylation was quantified by dividing intensity of NMP-nsp9 band with intensities of NMP-nsp9+free nsp9 bands from two independent replicates. (E) Two different orientations of the 5’-adenine (A1) in the base-up pose and their interactions with nsp12 residues in the RNA-nsp9:RTC complex (PDB ID 8GWE (magenta) and PDB ID 8SQK (dark grey)). (F) RNAylation activity of WT-RTC and RTC containing indicated mutants of nsp12 involved in RNA binding. The reaction products were analyzed as in Fig. 1D. %RNAylation was quantified by dividing intensity of RNA-nsp9 band with intensities of RNA-nsp9+free nsp9 bands from two independent replicates. Molecular graphics generated using ChimeraX ^46^. Data of replicates represented as mean and standard error of mean.

A wealth of information on the mechanism of these reactions performed by NiRAN has come from structural studies combined with mutational and biochemical analyses carried out in different laboratories around the globe^17,21,23,25–27^. Additionally, mechanistic studies of kinases, such as casein kinase 1, and pseudokinase Selenoprotein-O^28,29^, which have the same fold as NiRAN’s, have provided additional insights^17,30,31^. Structures of various complexes of SARS-CoV-2 RTC having RNA and/or nucleotide substrate/s, in addition to nsp9, bound to NiRAN have revealed that the substrates can occupy three different but overlapping binding poses within a single active site – base-in, base-out and base-up poses^23,32^ (Fig. 1B).

In the base-in pose, the base of the nucleotide is oriented towards the inside of NiRAN into a pocket that is specific for guanine, referred to as the G-pocket^23,27,33,34^. Guanine specificity is achieved through base-specific interactions made by residues nsp12-Arg55, nsp12-Tyr217 and nsp12-Thr120^23,27,33^. This nucleotide binding pose is similar to that adopted by ATP bound to kinases having NiRAN-like fold, such as casein kinase 1 that phosphorylates its protein substrates^17,28^. GDP bound to NiRAN in the base-in pose deNMPylates or deRNAylates nsp9 (Fig. 1B). Structural studies have shown that GTP or its analogues can also bind in this pose (Table S1). Recently, it was proposed that GTP bound in the base-in pose can be hydrolyzed to GDP by the NiRAN domain when RNAylated nsp9, but not NMPylated one, is bound^34^. The newly formed GDP then catalyzes deRNAylation/deNMPylation.

The other two nucleotide binding poses are associated with performing NMPylation and RNAylation reactions. In the base-out pose, the nucleotide is oriented in NiRAN for nucleotide transfer reaction such that it is rotated by almost 180° with respect to the base-in pose (Fig. 1B)^21,35,36^. The same binding mode is adopted by ATP bound to the bacterial enzyme SelO that catalyzes a reaction similar to that of NiRAN^28,29^. The base-out pose was initially proposed to be the nucleotide binding pose for NMPylation^17,21^. However, a shortcoming of this proposal has been that all the structures with nucleotide bound in the base-out pose are either ADP or GDP^21,26,35,36^, but none with the catalytically relevant triphosphate-containing NTP (Table S2). Recently, a structure of the pre-catalytic state of 5’-pppA-RNA bound to RTC and nsp9 was published which showed the orientation of the 5’ nucleotide of the RNA during RNAylation to be in base-out pose^37^.

In the base-up pose, the nucleobase is oriented towards the interface between NiRAN and RdRp, almost perpendicular to that of base-in and base-out poses (Fig. 1B, C). Structures of the post-catalytic RNAylated nsp9 bound to RTC show that the 5’-adenosine of the RNA is in the base-up pose (Table S3) ^27^. Some of these structures also have analogues of GTP and GDP bound in the base-in pose, which provide a snapshot of the base-up orientation of the 5’ nucleotide of the RNA during deRNAylation reaction ^23,27^. A recent structure of nsp9-bound RTC has UMPPCP, an analogue of UTP, bound in the base-up pose, proposed to be in the pre-catalytic state of NMPylation^23^. In this pose, even though the nucleotide is bound to NiRAN, the base makes important contacts with a loop from the RdRp domain of nsp12. The structures of NMP-nsp9 bound RTC (post-catalysis) have the nucleobase in this pose too^27^. Based on the similarities of the orientations of the nucleotides in these structures, it has been proposed that the NTP for NMPylation and the 5’-adenine nucleoside of the RNA for RNAylation are bound in the same base-up pose for the respective reactions to occur^23,32^ (Fig. 1B).

The structural information on the NiRAN and RdRp domains favor a base-up pose for the NMPylation and RNAylation reactions. However, this conclusion poses a conundrum of how the same pose functions as a pocket specific for adenine for RNAylation and a promiscuous pocket for either ATP, GTP, UTP or CTP as substrate for the NMPylation reaction. Also, the evolutionary related SelO, which catalyzes a similar reaction, binds ATP in the base-out pose. Consequently, there remain ambiguities of the exact binding poses of these substrates. Through this study, we investigate the significance of the base-up, base-out and base-in poses in the context of RNAylation, NMPylation and deNMPylation. We also delve into the molecular underpinning of adenine specificity of RNAylation. Furthermore, we demonstrate nucleotide transfer using the substrate GpppN, which has not been reported previously, and decipher its mode of interaction with the NiRAN domain. Finally, we also explore the conditions under which the RTC would perform RNAylation reaction preferentially over NMPylation, when the ligands for both are available, as will be the case in the cellular milieu.

## Results

### Base-up pose is the primary nucleotide binding mode during RNAylation but not during NMPylation

To identify which of the two poses -the base-up or the base-out - was important for nucleotide binding during NMPylation, we mutated residues that were involved in distinct interactions in the two poses. The NMPylation and RNAylation of pure nsp9 by wild type RTC (Fig. S1) and its mutants were quantified and compared. In the structure of UMPPCP bound to RTC:nsp9 in the base-up pose (PDB ID 8SQ9), nsp12-Asn39 from the NiRAN domain and nsp12-Asn713 from the NiRAN-associated palm motif (NPalm) of the RdRp are within hydrogen bonding distance of O4 and O2 of the uracil base, respectively^23^ (Fig. 1C). These residues also interact with the first adenine of 5’-pA-RNA-nsp9. These interactions have been proposed important for both NMPylation and RNAylation^23^. In contrast, the two residues do not interact with nucleotide bound in the base-out pose (PDB ID 7CYQ & 6XEZ), and consequently are expected to be of no significance to nucleotide binding in this pose. nsp12-N713A mutation did not affect NMPylation and mutation of nsp12-Asn39 to alanine also did not affect AMPylation, CMPylation or UMPylation activity of the NiRAN domain (Fig. 1D, S1, S2A-D). This indicated that the interactions made by nsp12-Asn39 and nsp12-Asn713 in the base-up position was not important for NMPylation.

Interestingly, GMPylation was severely affected by nsp12-N39A mutation (less than 15% of the wild type activity) (Fig. 1D, S2A, C). Since the mutant RTC could use ATP/CTP/UTP to NMPylate nsp9 very well, we did not think that GTP would be the only nucleotide that bound to NiRAN in the base-up pose during GMPylation. Like GMPylation, IMPylation of nsp9 using inosine triphosphate (ITP), an analogue that differs from GTP by lack of the amino group at the base’s C2 position and which could be NMPylated by RTC, was also affected by the mutation (Fig. S3A-D). Furthermore, modified pyrimidine-containing nucleotide – ^5Me^CTP and ^5Me^UTP – could also be used by RTC for NMPylation of nsp9, and these reactions were unaffected by nsp12-N39A mutation (Fig. S3A-D). The results suggested that the reduction in NMPylation by nsp12-N39A mutated RTC was specific to guanosine-like nucleotide that had an oxo-group at the sixth position of the purine ring. Though it is not obvious from the structure why the mutation affected only GMPylation/IMPylation, one possible reason could be that the GTP/ITP bound with high affinity, mediated by O6 of the purine, to a base-specific pocket created by the mutation, which prevented the nucleotide from binding in a reaction-competent pose.

The structures of post-catalytic RNAylated nsp9 bound to RTC show that the first nucleotide of 5’-pppA-RNA is proposed to be in the base-up pose ^23,27^. In these structures the first nucleotide of the RNA has been captured in the base-up pose bound to NiRAN via two binding modes (Fig. 1E; Table S3). In one set of structures, the adenine base of the 5’-nucleotide forms hydrogen bond with nsp12-Asn39 and nsp12-Asn713^27^ (PDB ID 8GWE). In the other set, the adenine base is flipped such that it does not interact with nsp12-Asn39, but interacts with nsp12-Asn713 and the neighboring nsp12-Asp711 (PDB ID 8SQK & 8SQJ) ^23^. Consequently, we expected the point mutants nsp12-N39A, nsp12-N713A and nsp12-D711A to affect RNAylation. While wild type RTC RNAylated 80% of nsp9 with a 5-mer RNA (5’-ppp-AUUAA), nsp12-N39A and nsp12-N713A mutants showed 0% and ∼15% product-formation, respectively (Fig. 1F, S4). This demonstrated the importance of the interactions made by nsp12-Asn39 and nsp12-Asn713 with the 5’-nucelotide during RNAylation reaction, which was consistent with the proposal made by Small et al., 2023^23^, that the 5’-nucleotide of the RNA takes the base-up pose. In contrast, RTC with nsp12-D711A mutation could RNAylate ∼50% of nsp9 indicating a less critical role of nsp12-Asp711 for the reaction (Fig. 1F, S4).

### nsp12-Asp711 contributes to the base specificity for RNAylation

As mentioned earlier, RNAylation occurs only if the 5’-nucleotide of the RNA is an ATP (5’-pppA-RNA) ^17^. We wondered if nsp12-Asn39, nsp12-Asp711 and/or nsp12-Asn713, which interact with the base of the 5’ adenine nucleotide of the RNA substrate, contributed to base specificity. For this, we checked if RTC with point mutation of the above residues could RNAylate nsp9 using 5’-pppG-RNA. To test this, we used a 10-mer 5’-pppG-RNA. While the wild type, nsp12-N39A and nsp12-N713A mutants showed no RNAylation using 5’-pppG-RNA as the substrate, RTC containing nsp12-D711A and nsp12-D711N mutations showed a weak RNAylation activity (Fig. 2A-B, S5). These mutations, thus, revealed the significance of nsp12-Asp711 in specific recognition of the 5’ nucleotide of the RNA.

**Figure 2:**
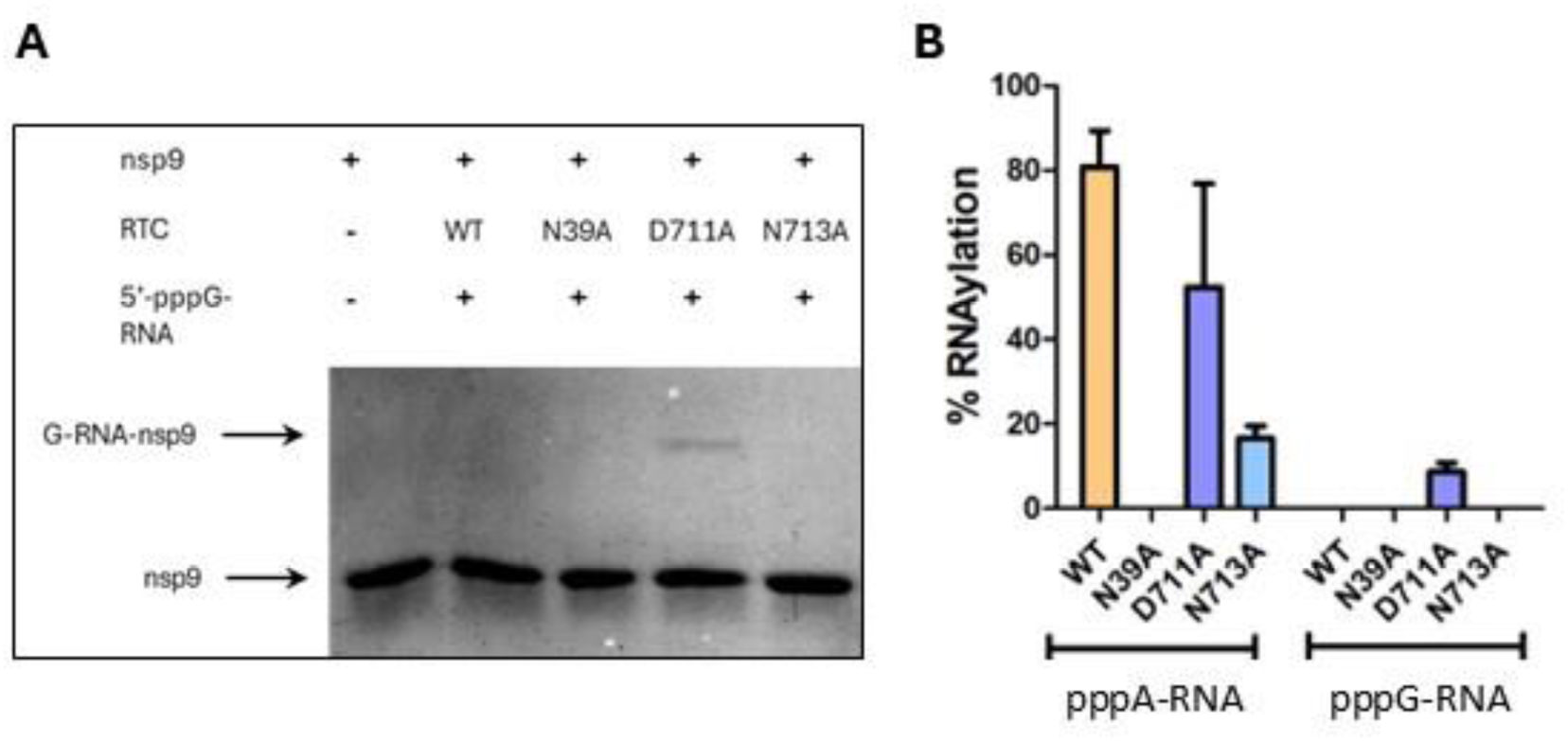
nsp12-Asp711 contributes to base-specificity for RNAylation. (A) A representative gel of two independent replicates showing RNAylation using 5’-pppG-RNA with WT RTC and RTC containing indicated mutants of nsp12. (B) RNAylation activity of WT-RTC and RTC containing indicated mutants of nsp12 using 5’-pppA-RNA (5mer) and 5’-pppG-RNA (10mer). Reaction products were analyzed as in Fig. 1F from two independent replicates.

As the nsp12-D711A could RNAylate both 5’-pppA-RNA and 5’-pppG-RNA, though only feebly (Fig. 2B, S4, S5), it broadened the substrate specificity of RTC for RNAylation of nsp9. Mutation of nsp12-Asp711 to alanine or to aspargine removed the negatively charged sidechain of nsp12-Asp711, which lies in the vicinity of the purine base. The side chain of nsp12-Asp711 contributed to the stable binding of 5’-pppA-RNA as it formed hydrogen bond with N6 of the 5’ adenine base. With a guanine at this position, the lone pair electrons of O6 of the base and the negatively charged side chain of nsp12-Asp711 would electrostatically repel each other preventing the binding of 5’-pppG-RNA. nsp12-D711A and nsp12-D711N removed the electrostatic repulsion allowing binding of 5’-pppG-RNA (Fig. S6). Interestingly, as shown previously, in comparison to wild type RTC, nsp12-D711A mutation reduced RNAylation using 5’-pppA-RNA by only about 30% (Fig. 2B, S4), highlighting that nsp12-Asp711 does not significantly contribute to substrate binding, but does play an important role in preventing 5’-pppG-RNA as an RNAylation substrate.

### Base-out pose is the primary nucleotide binding pose for NMPylation

Comparison of structures having nucleotide bound in the base-up and base-out poses revealed that the residue nsp12-Asn52, which is highly conserved in coronavirus nsp12 (Fig. 3A), was close to an oxygen atom of α-phosphate of the nucleotide in the base-out pose (∼4.2 Å; Fig. 3B). In the base-up position, the side chain of this residue was pointing away from the nucleotide exposed to the solvent and located in the vicinity of the backbone oxygen of nsp12-Arg74^23^ (PDB ID 8SQ9). In the homolog SelO, the residue equivalent to nsp12-Asn52 is Arg93. SelO-Arg93 stacks with the base of the ATP bound in the base-out pose and interacts with the β-phosphate (Fig. 3C). As the structures of SARS-CoV-2 RTC having nucleotide in the base-out pose have either ADP or GDP bound instead of an NTP (Fig. 3B, Table S2), we modelled a structure of RTC:nsp9 bound to an ATP using AlphaFold 3^38^ (Fig. 3D). In this model nsp12-Asn52 formed hydrogen bond with an oxygen atom of the α-phosphate. To find if this interaction was important for NMPylation of nsp9, we mutated nsp12-Asn52 to alanine and performed the NMPylation assay. The percentage activity of AMPylation, GMPylation and CMPylation shown by the nsp12-N52A mutant was less than 5% of the wild type activity, while UMPylation was about 15% (Fig. 3E, S7A-C). The significant decrease in NMPylation highlighted the importance of the interaction for the reaction. In turn, this implied that the base-out pose is an important nucleotide binding mode for NMPylation.

**Figure 3:**
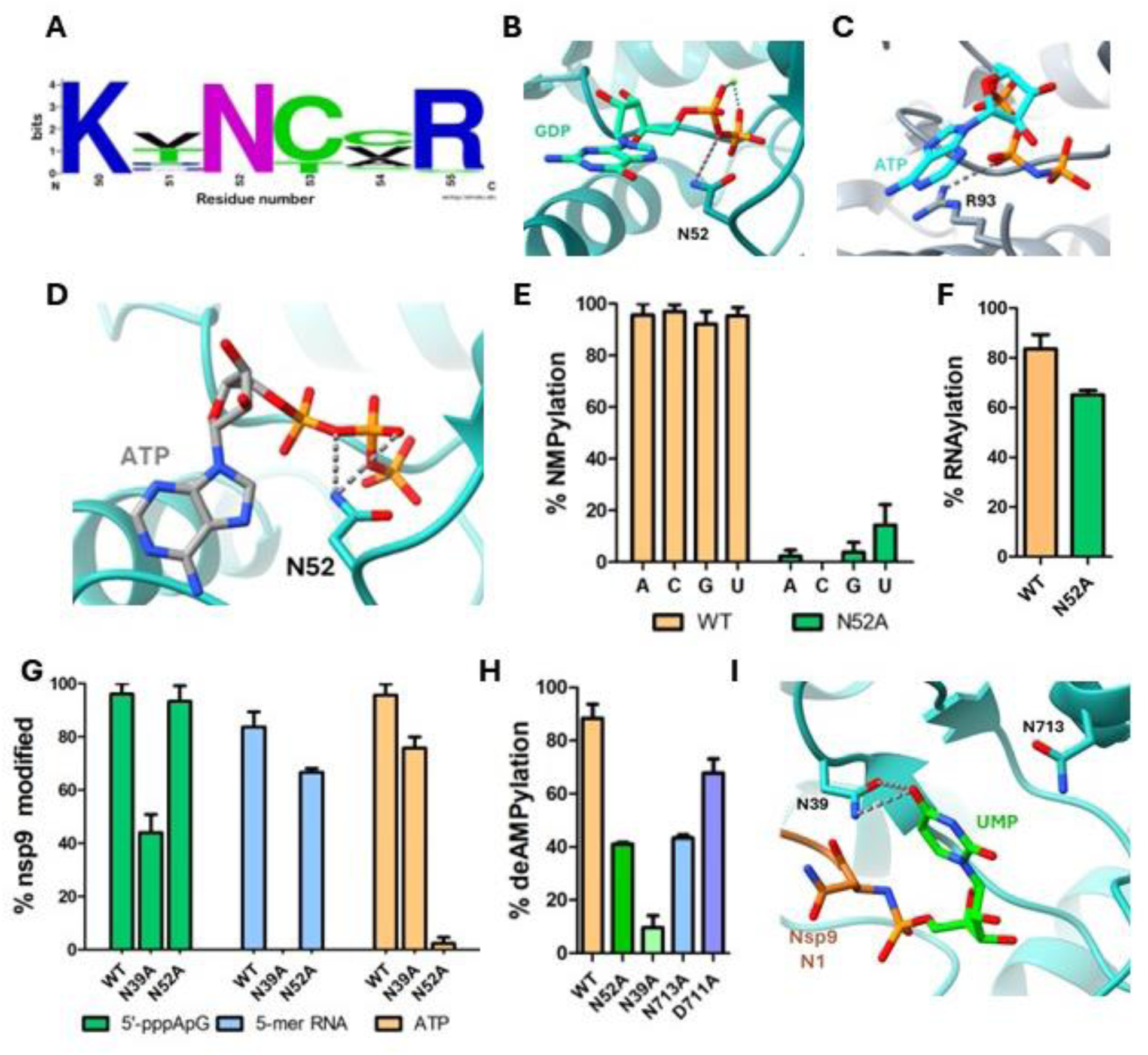
Base-out pose the nucleotide binding pose for NMPylation and base-up the nucleotide binding pose during deNMPylation. (A) Weblogo showing conservation of nsp12-Asn52. Generated from an MSA of 93 nidovirales nsp12 sequences using Clustal-Omega^47^. (B) Interactions of nsp12-Asn52 and GDP in base-out pose (PDB ID 7CYQ). (C) Interactions of SelO-Arg93 with ATP in Pseudokinasae SelO active site (PDB ID 6XEZ). This pose is similar to the base-out pose (B). (D) Interactions of nsp12-Asn52 and ATP in base-out pose in a model generated using Alphafold-3^38^. (E) NMPylation activity of WT-RTC and RTC containing nsp12-N52A mutation. Reaction products were analyzed as in Fig. 1D from two independent replicates. (F) RNAylation activity of WT-RTC and RTC containing nsp12-N52A mutation. Reaction products were analyzed as in Fig. 1F from two independent replicates. (G) Nucleotide transfer activity of WT-RTC and RTC containing indicated mutants of nsp12 using 5’-pppApG, 5’-pppA-RNA (5mer) and ATP. Reaction products were analyzed as in Fig. 1D for %AMPylation and Fig. 1F for %RNAylation from at least two independent replicates. (H) DeAMPylation activity of WT-RTC and RTC containing indicated mutants of nsp12. Reaction products were analyzed as in Fig. 1D from at least two independent replicates. %deAMPylation was quantified by dividing intensity of free nsp9 band with intensities of AMP-nsp9+free nsp9 bands from at least two independent replicates. (I) Interactions of UMP-nsp9 in the base-up pose (PDB ID 8GWO).

As proposed earlier^23^ and as inferred above, 5’-nucleotide of the RNA in an RNAylation reaction takes the base-up pose. We next tested if nsp12-N52A mutation affected RNAylation like it affected NMPylation. In our experiments we found that the mutated NiRAN could still RNAylate nsp9 (Fig. 3F, S8). This further confirmed that RNAylation happened with the 5’-adenine nucleotide positioned in the base-up pose. Using the nsp12-N52A mutant that specifically affects NMPylation and nsp12-N39A mutant that affects RNAylation, we probed the minimum length of an oligonucleotide that would take a binding pose akin to a long RNA substrate and RNAylate nsp9. We began with the shortest oligonucleotide – a dinucleotide with the sequence 5’-pppApG-3’, which satisfied the sequence requirements of RNAylation, i.e., having a triphosphate at the 5’-end and an adenine as the first base. As observed previously by Park *et al*, 2022^17^, we found that the RTC could use dinucleotide to modify nsp9. Interestingly, the mutant RTC containing nsp12-N52A mutation also transferred the dinucleotide to nsp9 as well as the wild type, whereas that having nsp12-N39A mutation had a considerably reduced transfer efficiency (Fig. 3G, S9). This result implied that during RNAylation of nsp9 the dinucleotide bound NiRAN in the base-up pose just like the first nucleotide of a longer RNA would do. Consequently, in some of the subsequent studies, we used the dinucleotide substrate instead of a longer 5’-pppA-RNA for studying RNAylation. A recent report proposed that nsp12-Asn52 and nsp12-His75 make important interactions with the 5’-adenine nucleotide of the RNA during RNAylation based on the structures of the pre-catalytic state of NiRAN in which the nucleotide was in the base-out pose^37^. We found that neither nsp12-N52A mutation nor the nsp12-H75A mutation affected RNAylation significantly. In our experiments, neither the nsp12-N52A mutation (Fig. 3F, S8) nor the nsp12-H75A mutation significantly affect RNAylation (Fig. S10A, B). These results proved that nsp12-Asn52 and nsp12-His75 did not play a significant role in RNAylation, and led us to conclude that the base-out pose is not relevant for RNAylation.

### Base-up pose is the primary nucleotide binding pose during deNMPylation

Through the mutation of nsp12-Asn52, we established that NMPylation happens primarily through the base-out pose, and that the residue does not have a significant role in catalyzing RNAylation. We next wanted to find if nsp12-Asn52 has a role in deNMPylation. DeNMPylation requires GDP to be bound in the base-in pose to the NiRAN’s G-pocket. Based on the structures of RTC having nucleotide bound in the base-in pose, one can categorize five kinds of interactions formed by nsp12-Asn52 with a guanine nucleotide in the base-in pose (Fig. S11; Table S1). Comparison of these structures showed that nsp12-Asn52 did not make a conserved interaction with the nucleotide in the base-in pose. To test if nsp12-Asn52 is important for deNMPylation, we mutated it to alanine and found that the mutant RTC could deAMPylate 40% of AMP-nsp9, while the wild type RTC could deAMPylate 90% (Fig. 3H, S12). Unlike NMPylation, the mutation did not fully inhibit deAMPylation.

In contrast, we noted that nsp12-N39A mutation reduced deAMPylation by almost 80%, while the NPalm mutants nsp12-N713A and nsp12-D711A reduced the activity by ∼45% and ∼20%, respectively (Fig. 3H, S12). The observation that nsp12-N39A mutation significantly affected deAMPylation, and nsp12-N713A to a lesser extent, despite not being part of either the G-pocket or the catalytic pocket led us to infer that the residues might interact with the nucleotide and stabilize the AMP-nsp9 complex for deAMPylation. This implied that the nucleotide covalently linked to nsp9 is oriented in the base-up pose and stabilized by nsp12-Asn39 and nsp12-N713A. The inference is consistent with the structure of a UMPylated nsp9 bound to RTC in which the UMP is in the base-up position and is within hydrogen bonding distance of the side chain of nsp12-Asn39 (Fig. 3I). Also, we think that the interaction between nsp12-Asn52 and GDP bound in the base-in pose is not as critical for deNMPylation, since the mutant RTC could deAMPylate almost 40% of AMP-nsp9.

### NiRAN can use GpppN to NMPylate nsp9

When carrying out deRNAylation and deAMPylation reactions, we observed that with increasing GDP concentration we consistently achieved near 100% deRNAylation of RNAylated nsp9, while deAMPylation was never above 80% (Fig. 4A-B, S13). As the reaction mix in the deRNAylation reaction did not have NTP, we wondered if the NiRAN domain was re-AMPylating nsp9 using the newly formed GpppA. To test this, we carried out NMPylation reaction of nsp9 by RTC using GpppA or GpppG. We found that the RTC could NMPylate nsp9 using either GpppA or GpppG (Fig. 4C-D, S14-18). Using mass spectrometric analysis, we found that when GpppA was used, only AMPylation of nsp9 was observed and there was no hint of GMPylation (Fig. 4E). Furthermore, AMPylation of nsp9 was not observed using ApppA (Fig. 4D, S15). From these observations we inferred that for the reaction to use NpppN as the substrate, at least one of the bases of the substrate has to be guanine. The inference led us to conclude that during the reaction the guanine base of GpppN binds to the highly specific G-pocket in the base-in pose.

**Figure 4:**
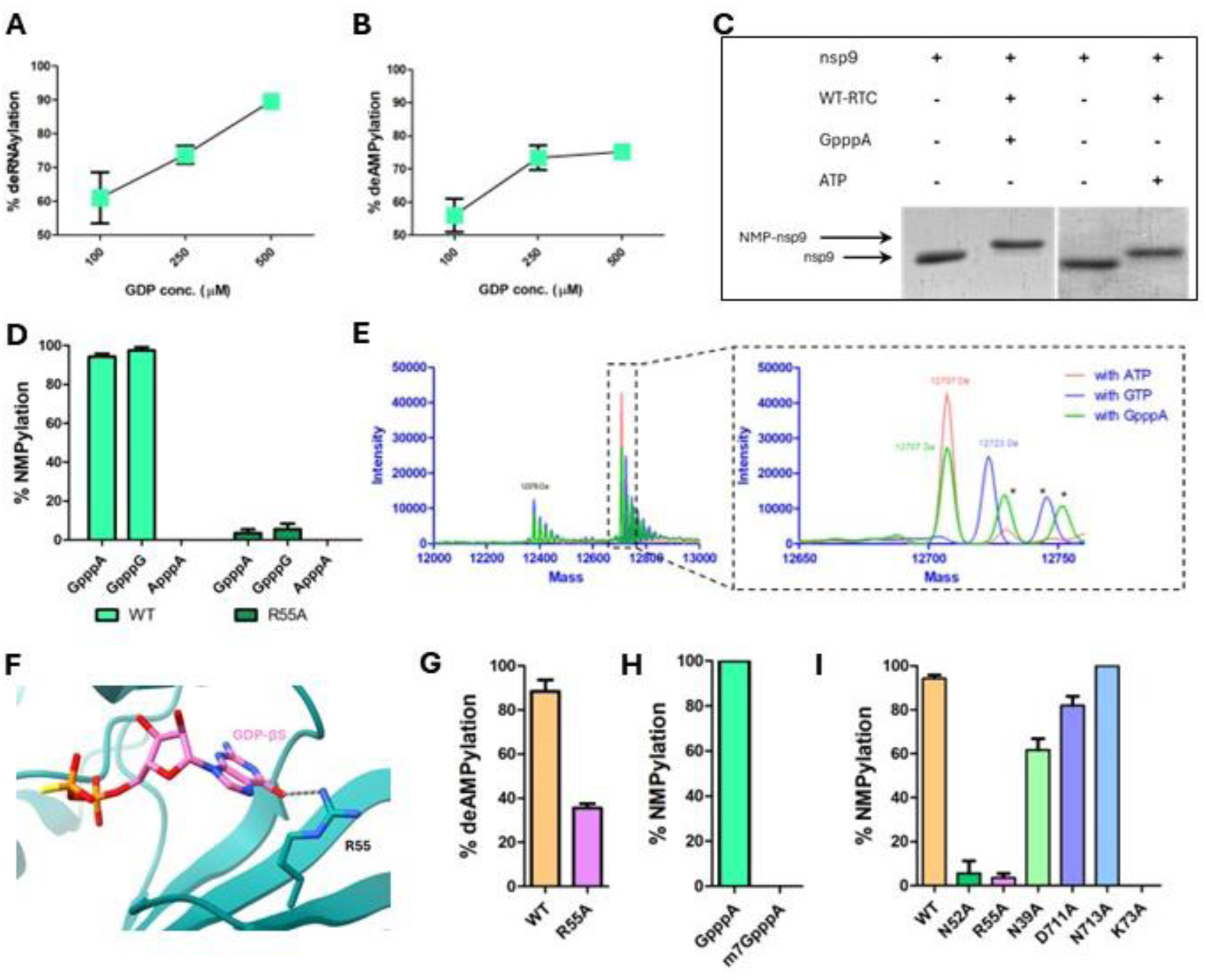
NiRAN can use GpppN to NMPylate nsp9. (A) DeRNAylation by WT-RTC with increasing GDP concentration. %deRNAylation was quantified by dividing intensity of free nsp9 band with intensities of RNA-nsp9+free nsp9 bands from three independent replicates. (B) DeAMPylation by WT-RTC with increasing GDP concentration. Reaction products were analyzed as in Fig. 3H from three independent replicates. (C) A representative gel of three independent experiments showing NMPylation using GpppA by WT-RTC in comparison to that using ATP. (D) NMPylation activity of WT-RTC and RTC containing nsp12-R55A mutation using GpppA, GpppG and ApppA. The reaction products were analyzed as in Fig. 1D from three independent replicates. (E) Deconvoluted and native MS spectra showing the mass shift corresponding to AMP-nsp9 and GMP-nsp9 after nsp9 NMPylation using ATP, GTP and GpppA. (inset) Zoomed to show peaks within 12650 Da to 12760 Da range revealing distinct peaks corresponding to AMP-nsp9 and GMP-nsp9. The observed masses were within 1 Da relative of the expected masses. Asterisks indicate adducts. A representative of three independent replicates. (F) Interactions of GDP-βS with nsp12-Arg55 in the base-in pose (PDB ID 8SQK). (G) DeAMPylation activity of WT-RTC and RTC containing nsp12-R55A mutation. Reaction products were analyzed as in Fig. 3H from at least two independent replicates. (H) NMPylation activity by WT-RTC using GpppA and ^m7^GpppA. Reaction products were analyzed as in Fig. 1D from three independent replicates. (I) NMPylation activity by WT-RTC and RTC containing indicated mutants of nsp12 using GpppA. Reaction products were analyzed as in Fig. 1D from three independent replicates.

nsp12-Arg55 present in the G-pocket plays a role in the specificity and binding of guanine in the base-in pose through its interaction with the base (Fig. 4F). Consequently, RTC containing the mutant nsp12-R55A showed only 35% deNMPylation activity, while the wild type RTC showed 85% activity (Fig. 4G, S19A-B). Consistent with the role of nsp12-Arg55 in the binding of guanine to the G-pocket, we found that the nsp12-R55A mutation prevented NMPylation of nsp9 using GpppA (Fig. 4D, S14). Also, NMPylation of nsp9 was not observed when we used ^m7^GpppA instead of GpppA (Fig. 4H, S18C-D). This, we concluded, was because the methylated guanine was prevented from binding to the G-pocket due to the steric clash between the methyl group and the side chain of nsp12-Arg55. These results confirmed our inference that the guanine base of GpppN bound to the G-pocket during NMPylation.

To find the pose adopted by pppN during the NMPylation of nsp9 using GpppN, we carried out the reaction using the nsp12-Asn39 and nsp12-Asn52 mutants of RTC, respectively. While nsp12-N39A mutant showed 60% NMPylation, nsp12-N52A mutant showed less than 10% activity (Fig. 4I, S17). This implied that the pppN in GpppN was in the base-out pose during NMPylation reaction (Fig. 6D). nsp12-Lys73 has been identified as the catalytic residue for RNAylation, NMPylation, deRNAylation and deNMPylation. Mutation of this residue affects these activities^17,23^ (Fig. S18, S20). We also found that nsp12-K73A failed to perform NMPylation using GpppN, confirming that the lysine was the catalytic residue for this reaction too (Fig. 4I, S18A, D).

### NiRAN preferentially catalyzes NMPylation over RNAylation, which is altered by nsp13

NiRAN domain has the dual ability to NMPylate or RNAylate nsp9. To find which of these two reactions would preferentially occur when substrates for both the reactions were present simultaneously in the reaction mix, we first carried out an *in vitro* reaction containing RTC, nsp9 and a mix of 200 μM NTP (50 μM each of ATP, GTP, CTP and UTP) and 50 μM of the dinucleotide RNA 5’-pppApG. The mix mimicked the milieu in the replication vesicle where NTP is expected to be at a higher molar concentration than 5’-pppA-RNA. Under this reaction condition, we found almost 100% of nsp9 to be NMPylated. Even under equimolar concentrations of 50 μM NTP (12.5 μM of each NTPs) and 50 μM 5’-pppApG, nsp9 was NMPylated with very little RNAylation observed (Fig. 5A, S21). The result highlighted the strong preference of NiRAN to NMPylate than RNAylate nsp9. This led us to a physiological conundrum as to how RNAylation would occur in the NTP rich milieu where the replicated RNA genome has to be capped for its translation. We wondered if other components in the replication vesicle affect the preference of RTC to perform NMPylation over RNAylation.

**Figure 5:**
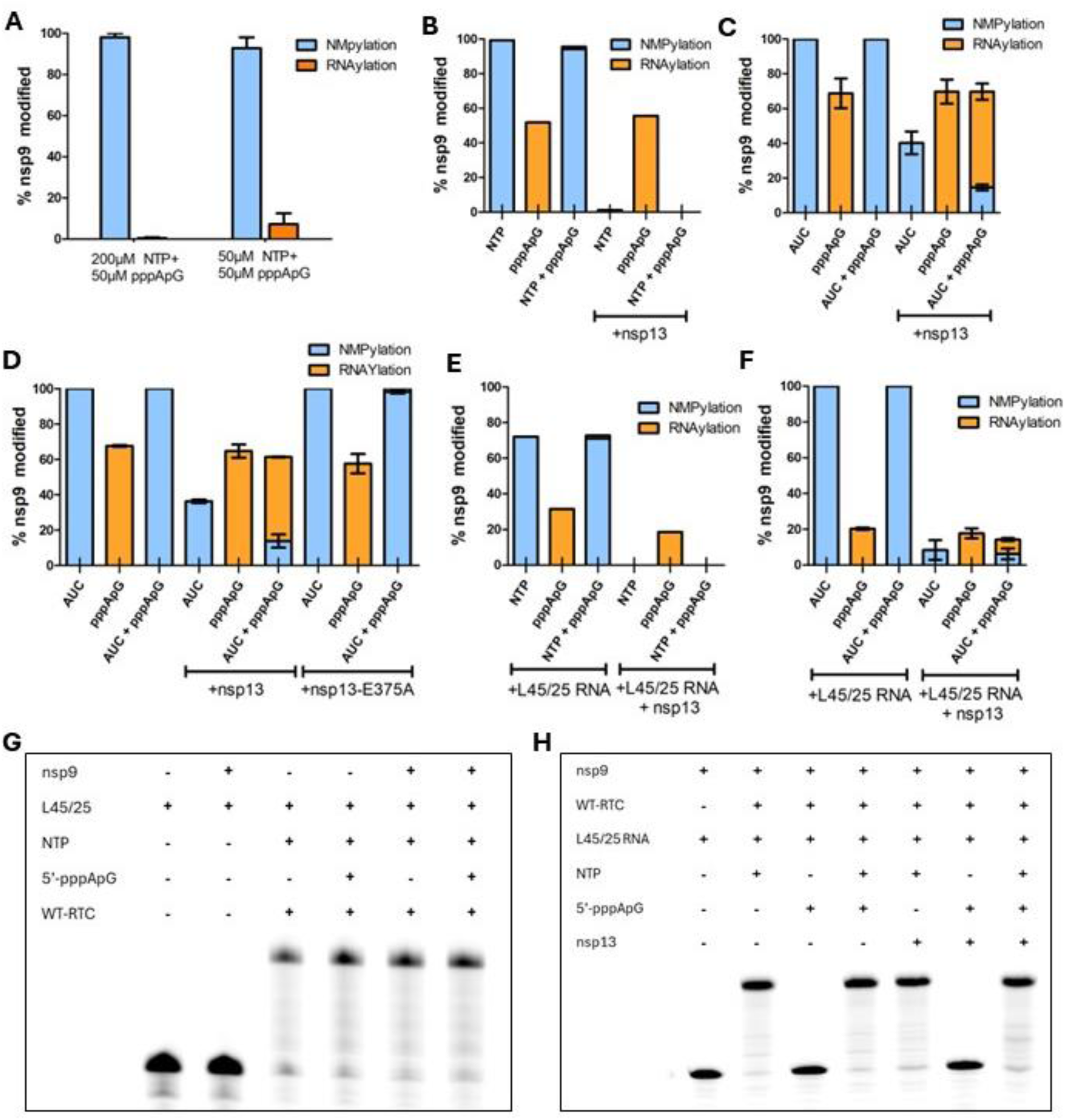
Regulation of NiRAN activities by nsp13 and RdRp. (A) Competition between NMPylation and RNAylation activities of WT-RTC using different ratios of NTP to 5’-pppApG. Reaction products were analyzed using SDS-PAGE and Coomassie staining and quantified using gel densitometry. %nsp9 modified was quantified by dividing the intensity of NMP-nsp9 band (for NMPylation) or RNA-nsp9 band (for RNAylation) the total intensity of RNA-nsp9/NMP-nsp9+free nsp9 bands from three independent replicates. (B) Competition between NMPylation and RNAylation activity of WT-RTC using 200 μM NTPs (ATP, GTP, CTP and UTP) and 50 μM 5’-pppApG, in the presence of RTC:nsp13. Reaction products were analyzed as in Fig. 5A from one replicate. (C) Competition between NMPylation and RNAylation activity of WT-RTC using 200 μM AUC (ATP+CTP+UTP) and 50 μM 5’-pppApG, in the presence of RTC:nsp13. Reaction products were analyzed as in Fig. 5A from three independent replicates. (D) Competition between NMPylation and RNAylation activities of WT-RTC using 200 μM AUC and 50 μM 5’-pppApG in the presence of WT-nsp13 and nsp13-E375A mutant. Reaction products were analyzed as in Fig. 5A from two independent replicates. (E) Competition between NMPylation and RNAylation activities of WT-RTC using 200 μM NTP and 50 μM 5’-pppApG, in the presence of L45/25 RNA and nsp13. Reaction products were analyzed as in Fig. 5A from 1 replicate. (F) Competition between NMPylation and RNAylation activities of WT-RTC using 200 μM AUC and 50 μM 5’-pppApG, in the presence of L45/25 RNA and nsp13. Reaction products were analyzed as in Fig. 5A from three independent replicates. (G) Polymerase activity of WT-RTC in the presence and absence of nsp9 using only 200 μM NTPs, or 200 μM NTPs and 50 μM 5’-pppApG. Reaction products were analyzed using Urea-PAGE and the bands were detected using fluorescently labelled RNA. A representative of two independent replicates. (H) Polymerase activity of WT-RTC using 200 μM AUC and 50 μM 5’-pppApG, in the presence and absence of nsp13. Reaction products were analyzed using Urea-PAGE and the bands were detected using fluorescently labelled RNA. A representative of three independent replicates. Plots B -F show percentage of RNAylation stacked over percentage of NMPylation for each reaction.

One of the components present during replication is nsp13, which has the ability to hydrolyse NTP to NDP^23,39,40^. Though there are some reports of SARS-CoV nsp13 hydrolysing the 5’-triphosphate of an RNA to 5’-diphosphate^39,41^, SARS-CoV-2 nsp13 did not hydrolyse 5’-pppApG in our experiments (Fig. S22). On addition of just 30 nM nsp13 to the above reaction mix containing 200 μM NTP and 50 μM 5’-pppApG, neither NMPylation nor RNAylation was observed (Fig. 5B, S23). We thought that our inability to observe NMPylated or RNAylated nsp9 was because the GDP generated from the nsp13-mediated hydrolysis of GTP was causing deNMPylaton and/or deRNAylation, as was observed previously^23^. We repeated the experiment again, but now we used a mix of only ATP, CTP and UTP with no GTP, referred to here as AUC mix. Using AUC, we found that only 15% of nsp9 were NMPylated, while almost 60% of nsp9 was RNAylated (Fig. 5C, S24). This result clearly demonstrated that nucleotide hydrolysis by nsp13 can tilt the reaction balance to favor RNAylation over NMPylation. Accordingly, replacement of nsp13 with its NTPase inactive Walker B mutant nsp13-E375A failed to prevent the strong preference of NiRAN to perform NMPylation (Fig. 5D, S25).

### RdRp catalyzed polymerization reaction decreases RNAylation of nsp9 by NiRAN but not vice-versa

All the reactions described so far were performed in the absence of an apo-RdRp domain. We wondered if the reactions carried out by NiRAN would be modulated by the binding of the corresponding RNA substrate to RdRp. Equally importantly, we wondered if the NTPase activity of nsp13 would deplete the NTP pool and hinder replication elongation along with NMPylation. To probe this, we performed NMPylation and RNAylation reactions in presence of an elongation substrate of RdRp made of a 45-mer template RNA annealed to a complementary 25-mer primer RNA, which we refer to as L45/25 (Fig. S26A). Upon addition of L45/25, we found that RNAylation reduced (Fig. 5B, E, S23, S27). We also tested if NMPylation or RNAylation modulated the elongation of L45/25 by RdRp. On comparison of the elongation product of RTC formed in the presence and absence of nsp9 we found that the polymerization reaction was unaffected when substrates for both NMPylation and RNAylation (5’-pppApG) were present in the reaction mix (Fig. 5G).

We next carried out the same comparative reactions in presence of nsp13. Earlier, we had shown that addition of nsp13 hindered NMPylation of nsp9 because it hydrolyzed NTP. We had also found that the hydrolysis of the nucleotide by nsp13 generated GDP that deNMPylated nsp9 and prevented us from quantifying the amount of NMPylated product formed. In presence of elongating L45/25 catalyzed by RdRp, we noticed the same effect, which implied that deNMPylation or deRNAylation were unaffected by the simultaneous polymerization reaction (Fig. 5E, S27). Nevertheless, to avoid the effects due to the formation of GDP, we used the AUC mix in the above reaction performed in presence of L45/25. As shown above, addition of nsp13 in the presence of AUC and 5’-pppApG reduced NMPylation and allowed RNAylation of ∼50% of nsp9 to occur. However, on addition of L45/25 RNA, only about 10% of nsp9 were RNAylated (Fig. 5F, S28). Interestingly, the elongation of L45/25 was unaffected by the presence of nsp13 (Fig. 5H; Fig. S26B). Interestingly, the same reaction carried out in the absence of the nucleotide mix AUC, which allowed RdRp to bind to L45/25 but not elongate its primer strand, also showed a lowering of RNAylated nsp9 by about 20% (Fig 5F, S28). This indicated that not only the polymerization of the RdRp substrate but just its binding reduced the efficiency of RNAylation.

## Discussion

The acknowledged essentiality of NiRAN domain for the lifecycle of nidoviruses makes it a prime target for therapeutics^27,42,43^. Though made primarily of a single domain, NiRAN can perform four different reactions involving distinct substrates, including NMPylation or RNAylation of the N-terminus of nsp9 or the deNMPylation or deRNAylation of NMPylated/RNAylated nsp9. Hence, the varied reactions carried out by NiRAN domain are excellent targets for therapeutics^27,42,43^. To develop effective inhibitors targeting these varied reactions, it is important to understand the binding poses of the respective substrates/ligands to NiRAN accurately. Interestingly, structures of NiRAN bound to its various substrates have revealed distinctly different binding modes even for the same substrate.

Based on the orientation of the nucleotide and the location of its base (first nucleotide in case of RNA), the modes have been categorized as base-up, base-out and base-in pose. The base-in pose, which can only be taken up by a guanine nucleotide due to the base specific interactions made by NiRAN residues in this pose, unambiguously represents the pose taken up by the capping GDP that deNMPylates/deRNAylates nsp9. However, the significance of the base-up and base-out pose for the reactions catalyzed by the NiRAN is ambiguous. Based on the similarities in the orientation of an NTP in the base-up pose and the first nucleotide of RNAylated nsp9 bound to NiRAN, it was proposed that the base-up pose is the catalytically relevant pose for both NMPylation and RNAylation, and the base-out pose was concluded to be insignificant^23,32^.

In the base-up pose the nucleotide interacts with the NiRAN residue nsp12-Asn39 and the NPalm residues nsp12-Asp711 and nsp12-Asn713. Mutation of either nsp12-Asn39 or nsp12-Asn713 affected RNAylation but not NMPylation. Mutation of nsp12-Asp711 did not significantly affect either RNAylation or NMPylation. In contrast, mutation of nsp12-Asn52, which interacts with the nucleotide only in the base-out pose and does not interact with the nucleotide in the base-up pose, is critical for NMPylation but not for RNAylation. Based on these observations, we concluded that NTP would be bound in the base-out pose during NMPylation, while the first nucleotide of an RNA substrate would be in the base-up pose during RNAylation (Fig. 6A, B). The binding of the NTP in the base-out pose is also similar to the orientation of the ATP bound to the evolutionarily related and structurally homologous pseudokinase SelO that AMPylates its protein substrates.

**Figure 6:**
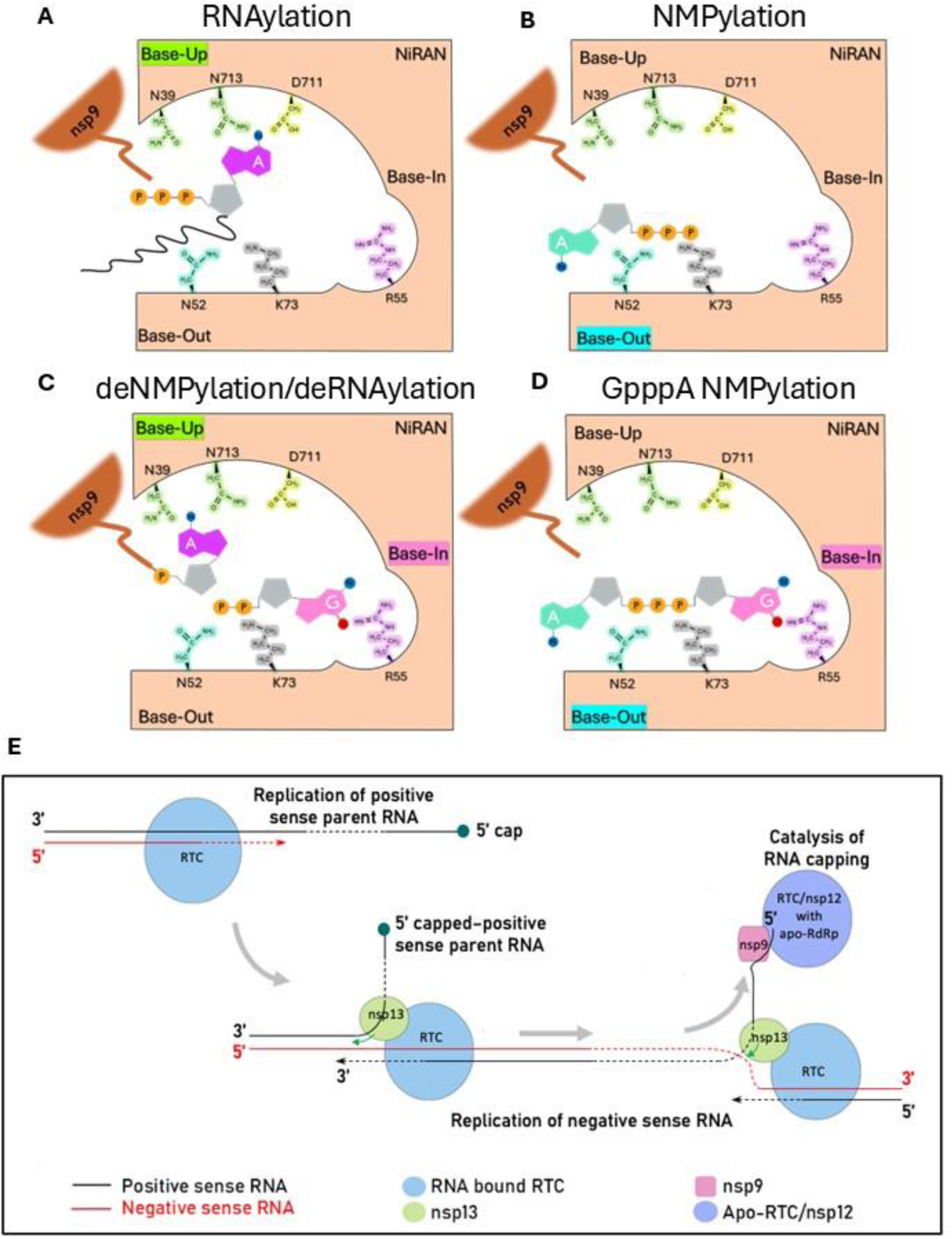
Schematic of substrate binding and regulation of NiRAN activities. (A) Schematic of RNA binding in base-up pose to NiRAN for RNAylation. The first adenine of the 5’-pppA-RNA is shown in magenta. (B) Schematic of ATP binding in base-out pose for NMPylation. Adenine is shown in cyan. (C) Schematic of GDP (pink) binding in base-in pose for deNMPylation/deRNAylation. First adenine base of RNA shown in magenta. (D) Schematic of GpppA binding to NiRAN during NMPylation where guanine (pink) binds to the G-pocket in the base-in pose and adenine (cyan) binds in the base-out pose. (E) Schematic showing initial rounds of RNA replication by RTC and subsequent RNA capping by RTC/nsp12 having apo-RdRp.

Surprisingly, nsp12-D711A facilitated feeble RNAylation of nsp9 using 5’-pppG-RNA, which neither the wild type nor any other mutant of nsp12 could. Based on this observation, we propose that nsp12-Asp711 blocks the binding of 5’-pppG-RNA via electrostatic repulsion between its side chain and the guanine base, but allows the binding of 5’-pppA-RNA for RNAylation reaction (Fig. S6). We also found that nsp12-N39A mutation affected deNMPylation while nsp12-N52A did not. Hence, we concluded that the orientation of the nucleotide of the NMPylated nsp9 is in the base-up pose. This conclusion is supported by the structure of UMP-nsp9 bound to RTC^27^, in which the nucleotide is in the base-up pose. Available structures show that, similar to NMPylated nsp9, the first nucleotide of RNA-nsp9 bound to RTC also occupies the base-up pose and, in accordance, we inferred that both deNMPylation and deRNAylation follow a similar mechanism.

In this study, we also discovered the ability of NiRAN to NMPylate nsp9 using GpppN, which was unknown previously. For this reaction to happen, we demonstrated that one of the two bases of this substrate has to be guanine. This led us to conclude that the substrate binding requires the guanine to bind to the G-pocket, and consequently mutation of nsp12-Arg55 in the G-pocket prevented NMPylation using GpppN. Also, nsp12-N52A hindered this reaction while nsp12-N39A, nsp12-N713A or nsp12-D711A did not. Hence, we concluded that during the NMPylation reaction GpppN is bound such that the guanine anchors the substrate to the G-pocket, while the remaining part (pppN) is bound in the base-out pose (Fig. 6D). While GpppN can be a substrate for the NMPylation reaction, methylation of the guanine base at the N7 position prevented NMPylation as the modified guanine cannot bind to the G-pocket. Methylation of guanine at the N7 position is catalyzed by nsp14 post formation of the core cap GpppA-RNA. Like GpppA-RNA, GpppA is also modified by nsp14^17^. As GpppN will be methylated soon after its formation, we think that NMPylation using GpppN may not be physiologically significant in the presence of nsp14. However, the finding provides new druggable geometries for developing anti-viral molecules.

In a reaction mix containing both NTP and 5’-pppA-RNA, we found NMPylation of nsp9 to be catalytically more favored over RNAylation. However, RNAylation becomes prominent over NMPylation when the SARS-CoV-2 helicase/NTPase nsp13, which can hydrolyze NTP^39,40^, is added to the reaction mix. We think that the hydrolysis by nsp13 reduced the concentration of NTP in the reaction mix, thus affecting the kinetics of NMPylation and facilitating RNAylation. Though it has been postulated that NMPylation of nsp9 could protect it from degradation in the host^23^ or that the NMPylated nsp9 could play a role in priming the replication of the viral genomic RNA^11,16,22–24^, physiological relevance of NMPylation of nsp9 remains unproven. NMPylation of nsp9 may stabilize the protein in the early stages of ORF translation. However, during later stages of translation, when nsp13 and other nsps downstream of it are synthesized, NMPylation of nsp9 would be stalled due to the NTPase activity of nsp13 and the existing NMPylated nsp9 would be deNMPylated by the nsp13 generated GDP. Interestingly, nsp12-Asn52, which is critical for NMPylation, is highly conserved in nidoviral nsp12. We propose that future studies of the lifecycle of a SARS-CoV-2 virus carrying the nsp12-N52A mutation will provide important insights on the physiological relevance of NMPylation.

Unlike NMPylation, the decrease in NTP concentration, due to their hydrolysis by nsp13, did not affect the RNA polymerization reaction catalyzed by the nsp12’s RdRp domain. Furthermore, we found that the binding of the RNA substrate to RdRp or its elongation by the polymerization reaction affected RNAylation of nsp9 considerably when concomitant substrates were present. These observations led us to conclude that the NiRAN domain is inefficient in RNAylation when the RTC has a substrate-bound holo-RdRp. However, NiRAN of an RTC having an apo-RdRp RNAylated nsp9 efficiently (Fig. 6E). Studies have also shown that just nsp12 with an apo-RdRp can also RNAylate nsp9^17,20,44^. To conclude, we propose a model for the regulation of the activities of NiRAN towards capping the 5’-end of a newly synthesized positive RNA strand. In the beginning, the parent genome is a capped positive sense RNA, which is replicated into a negative sense RNA by the RTC. In the next round of replication, RTC in complex with nsp13 unwinds the double-stranded RNA and uses the negative-strand RNA as template for synthesis of a new positive-strand^26,45^. The unwound positive-strand is capped by nsp9 bound to apo-RTC/nsp12 (Fig. 6E). In summary, the structure-based mutagenesis and reconstitution study reported here provides new insights and clarity to the mechanism of the steps that lead to formation of the core cap structure of the RNA by the versatile catalyst NiRAN.

## Acknowledgements

A.U. would like to thank the Indian Institute of Science Education and Research, Pune, for a PhD fellowship. P.S.S. would like to thank Department of Science and Technology (DST), Government of India, for Kishore Vaigyanik Protsahan Yojana fellowship. U.B. would like to thank the Department of Biotechnology (DBT), Government of India, for PhD fellowship. K.S. would like to thank Rotary Pune Sports City’s CSR initiative and MoE-STARS, Ministry of Education, Government of India, for funds for execution of this project. We would like to thank Dr. Sagar Pandit for providing the plasmid for T7 RNA polymerase. We would like to thank Mass Spectrometry facility at Indian Institute of Science Education and Research, Pune for MS data acquisition and Mass Spectrometry and proteomics core facility at National Centre for Cell Science, Pune for MS data analysis.

## Author contributions

A.U.: Methodology, investigation, data analysis and curation, validation, writing original draft, visualization. M.N.: Methodology, investigation, data analysis and curation, validation, reviewing of manuscript draft. P.S.S: Investigation, data analysis and curation, validation, reviewing of manuscript draft. U.B.: Investigation, data analysis and curation, validation, reviewing of manuscript draft. K.S.: Conceptualization, methodology, validation, writing original draft, supervision, project administration, fund acquisition.

