## Supplementary Information for "Mechanism of substrate binding by the SARS-CoV-2 NiRAN domain and modulation of its activities during replication"

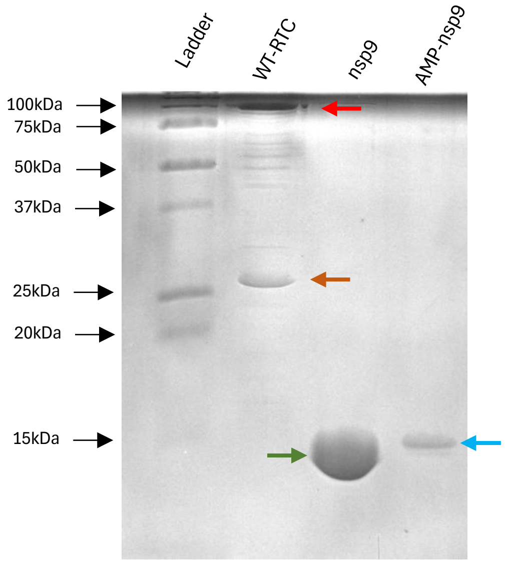


**Figure S1**: SDS-PAGE gel image of purified WT-RTC, nsp9 and AMP-nsp9. Protein bands of interest are marked using arrows – red for nsp12, brown for nsp8, green for nsp9 and blue for NMP-nsp9. nsp7 is not visible in the WT-RTC lane possibly because it is stained poorly. Bands between nsp12 and nsp8 in WT-RTC lane are possibly degradation products of nsp12, which has been observed by others too[1,2,3,4]. Gels are cropped to show only lanes of interest. The image is a representative of three independent replicates.

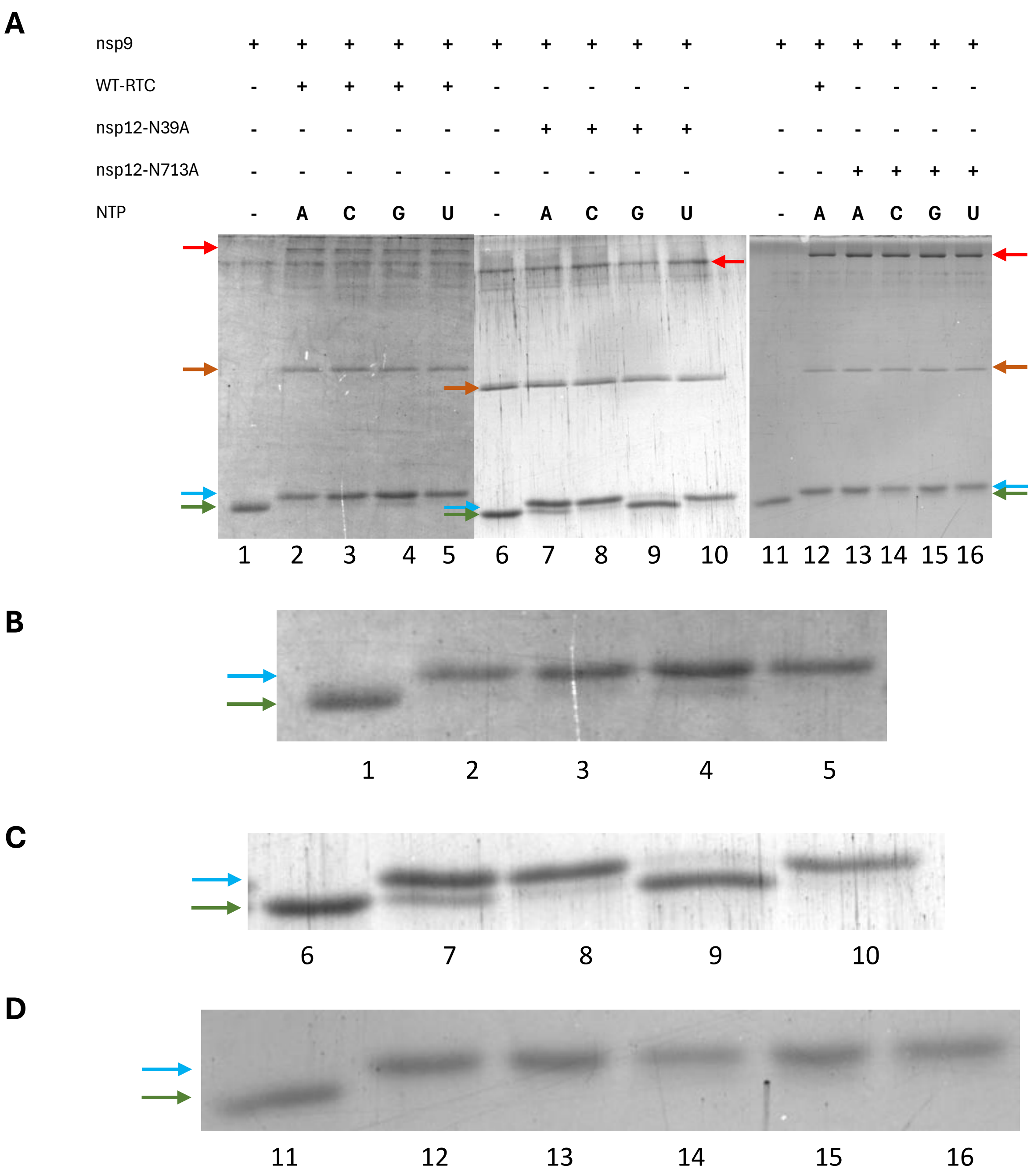


**Figure S2**: (A) Representative gel corresponding to Fig. 1D. Cropped SDS-PAGE gel images for nsp9 NMPylation experiment done with the WT-RTC and RTC containing indicated mutants of nsp12 using different nucleotide substrates shown along with no nucleotide control. Gels are cropped to show only lanes of interest. (B-D) Magnified and cropped image of the gels to highlight the modifications to nsp9. The numbers below the magnified images are lane numbers as in (A). All the images are representatives of three independent replicates.

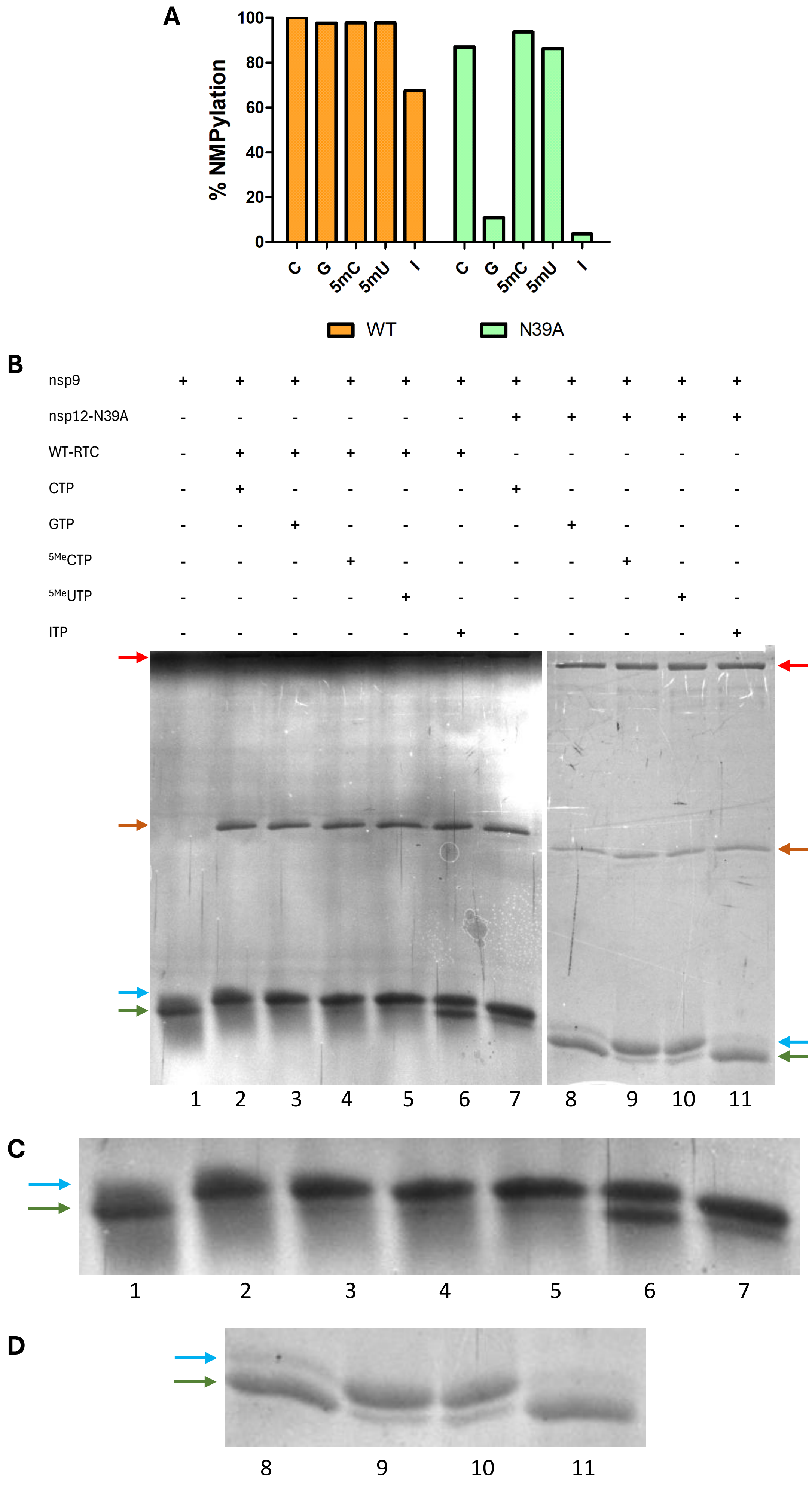
A
% NMPylation
100
80
60
40
20
0
C
G
5mC
5mU
I
C
G
5mC
5mU
I
WT
N39A
B
nsp9
nsp12-N39A
WT-RTC
CTP
GTP
5MeCTP
5MeUTP
ITP
1
2
3
4
5
6
7
8
9
10
11
C
1
2
3
4
5
6
7
D
8
9
10
11

**Figure S3:** (A) NMPylation activity by WT-RTC and RTC containing indicated mutants of nsp12 using ITP and other nucleotide substrates. Reaction products were analyzed as in fig. 1D from one replicate. (B) Representative cropped SDS-PAGE gel image for nsp9 NMPylation experiment carried out with WT-RTC and RTC containing indicated mutants using ITP and other nucleotide substrates. Different protein bands of interest are marked using arrows – red for nsp12, brown for nsp8, green for nsp9 and blue for NMP-nsp9. (C-D) Magnified and cropped image of the gels to highlight the modifications to nsp9. The numbers below the magnified images are lane numbers as in (B). Images are representatives of one replicate.

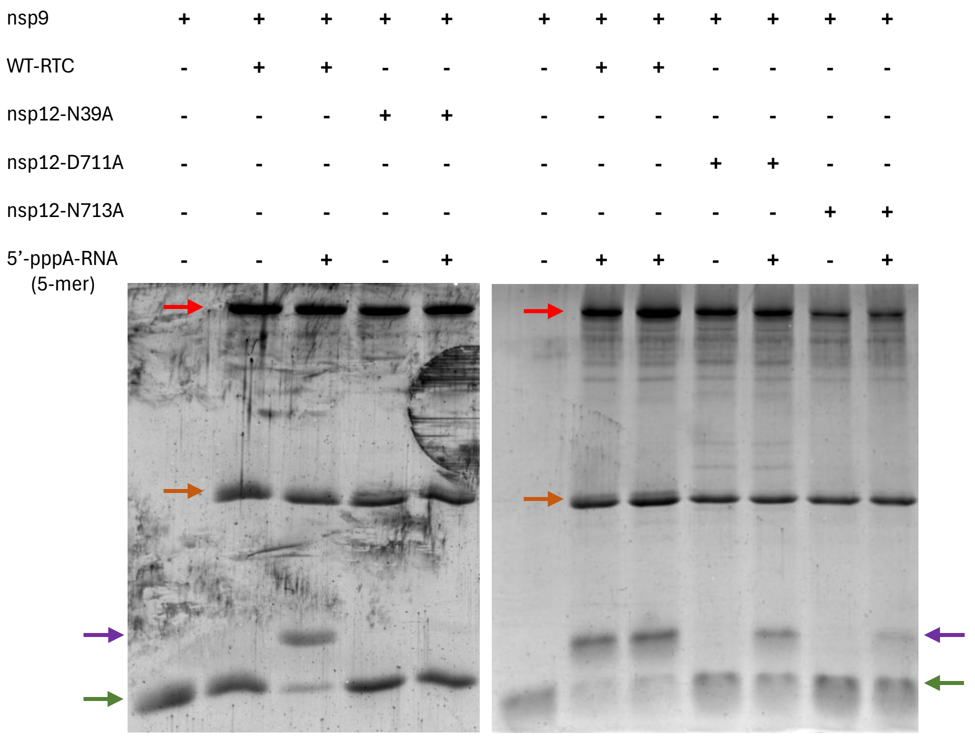


**Figure S4:** Representative gel corresponding to fig. 1F and 2B (pppA-RNA). Cropped SDS-PAGE gel images for nsp9 RNAylation experiment done with the WT-RTC and RTC containing indicated mutants of nsp12 using 5'-pppA-RNA shown along with no RNA control. Different protein bands of interest are marked using arrows – red for nsp12, brown for nsp8, green for nsp9 and purple for 5'-pA-RNA-nsp9. Gels are cropped to show only lanes of interest. All the images are representatives of three independent replicates.

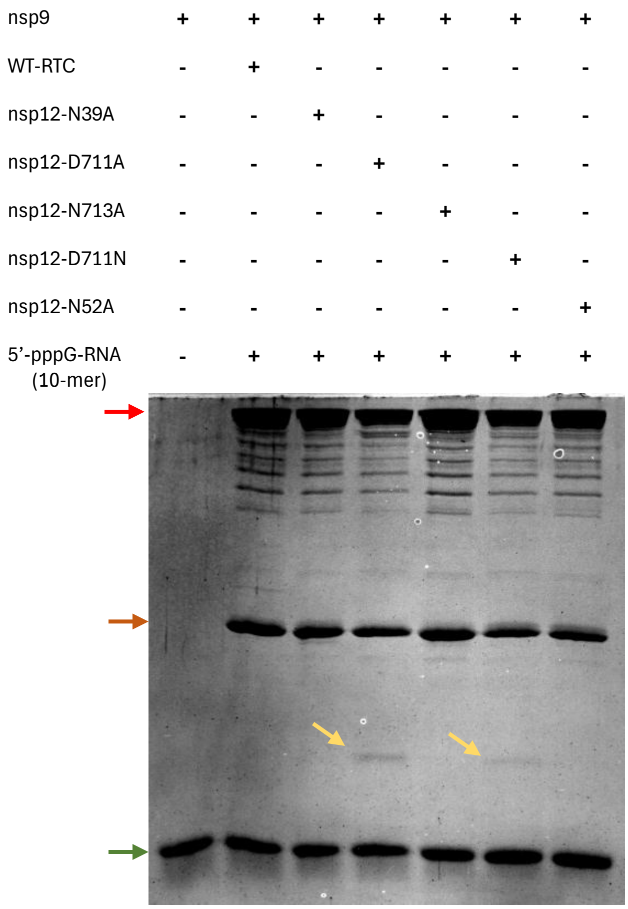


**Figure S5:** Representative gel corresponding to fig. 2A. Cropped SDS-PAGE gel images for nsp9 RNAylation experiment done with the WT-RTC and RTC containing indicated mutants of nsp12 which shows RNAylation using 5’-pppG-RNA by nsp12-D711A and nsp12-D711N mutants. Different protein bands of interest are marked using arrows – red for nsp12, brown for nsp8, green for nsp9 and yellow for 5’-pG-RNA-nsp9. Gels are cropped to show only lanes of interest. Image is representative of two independent replicates.

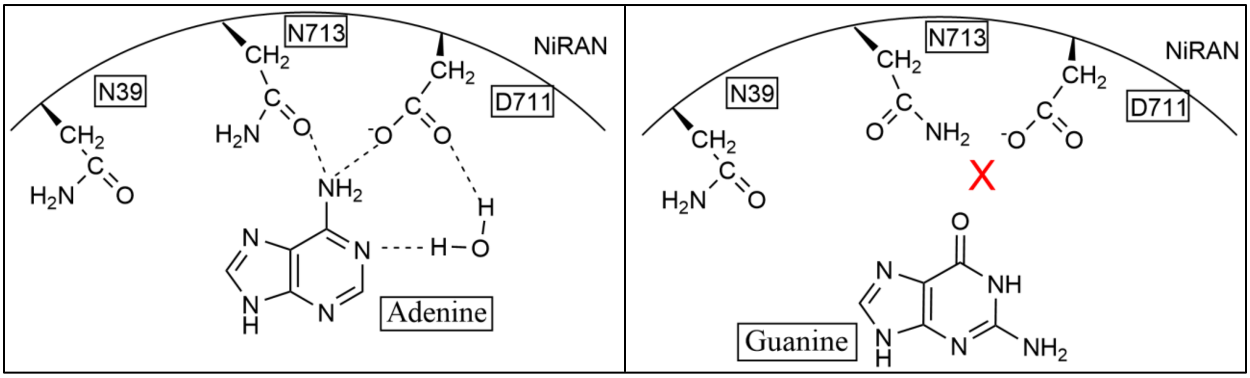


**Figure S6:** Schematic showing interactions made by NiRAN residues in base-up pose with an adenine base and guanine base. The red cross indicates the electrostatic repulsion between the carboxyl group of nsp12-Asp711 and the O6 of the guanine base.

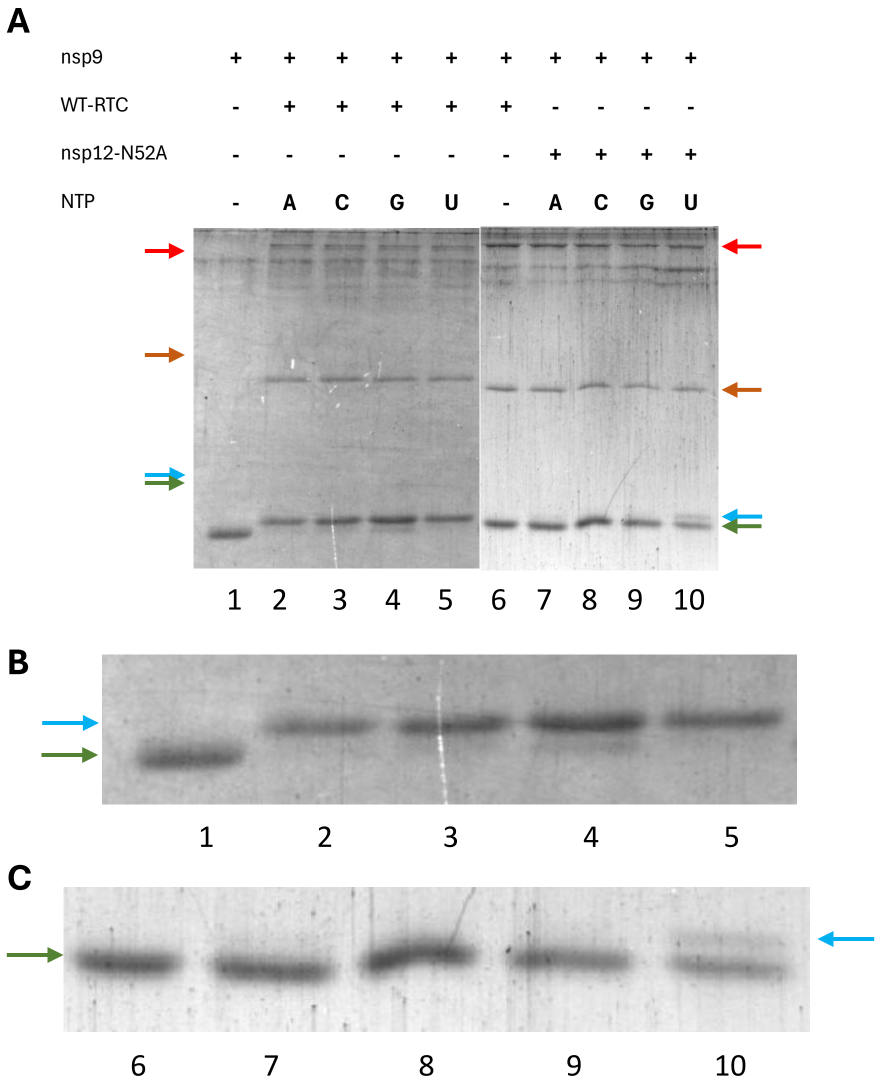


**Figure S7:** (A) Representative gel corresponding to fig. 3E. Cropped SDS-PAGE gel images for nsp9 NMPylation experiment done with the WT-RTC and RTC containing indicated mutants of nsp12 using different nucleotide substrates shown along with no nucleotide control. Different protein bands of interest are marked using arrows – red for nsp12, brown for nsp8, green for nsp9 and blue for NMP-nsp9. Gels are cropped to show only lanes of interest. (B-C) Magnified and cropped image of the gels to highlight the modifications to nsp9. The numbers below the magnified images are lane numbers as in (A). Images are representatives of three independent replicates.

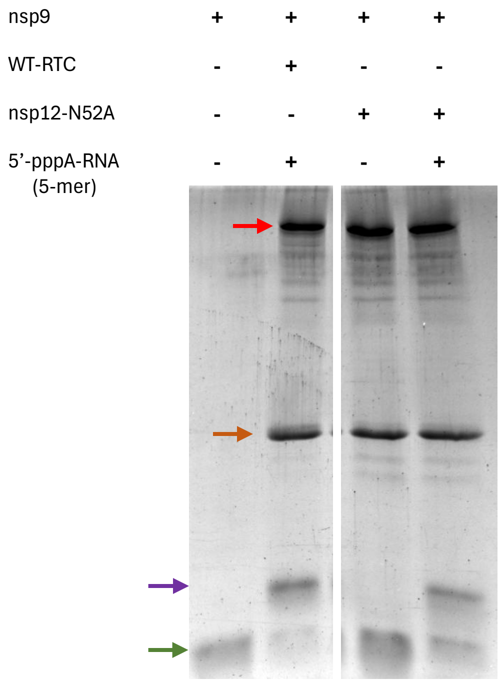


**Figure S8:** Representative gel corresponding to fig. 3F. Cropped SDS-PAGE gel images for nsp9 RNAylation experiment done with the WT-RTC and RTC having nsp12-N52A mutation using 5'-pppA-RNA shown along with no RNA control. Different protein bands of interest are marked using arrows – red for nsp12, brown for nsp8, green for nsp9 and purple for 5'-pA-RNA-nsp9. Gels are cropped to show only lanes of interest. Images are representatives of three independent replicates.

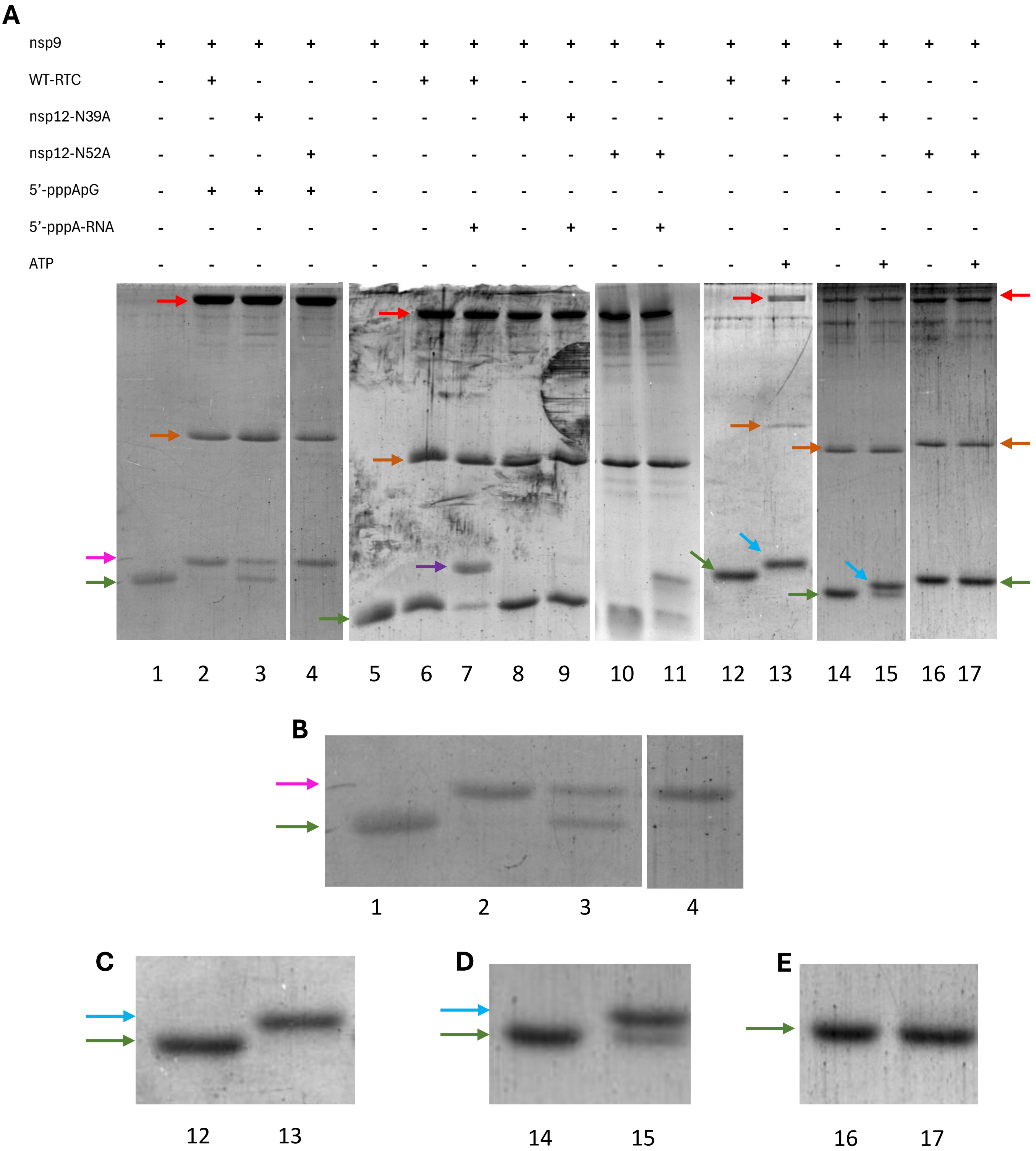


**Figure S9:** (A) Representative gel corresponding to fig. 3G. Cropped SDS-PAGE gel images for comparison of nsp9 RNAylation using 5’-pppApG and 5’-pppA-RNA and NMPylation using ATP experiment done with the WT-RTC and RTC containing indicated mutants of nsp12 shown along with no RNA control. Different protein bands of interest are marked using arrows – red for nsp12, brown for nsp8, green for nsp9, blue for AMP-nsp9, pink for 5’-pApG-nsp9 and purple for 5’-pRNA-nsp9. Gels are cropped to show only lanes of interest. (B-E) Magnified and cropped image of the gels to highlight the modifications to nsp9. The numbers below the magnified images are lane numbers as in (A). Images are representatives of at least two independent replicates.

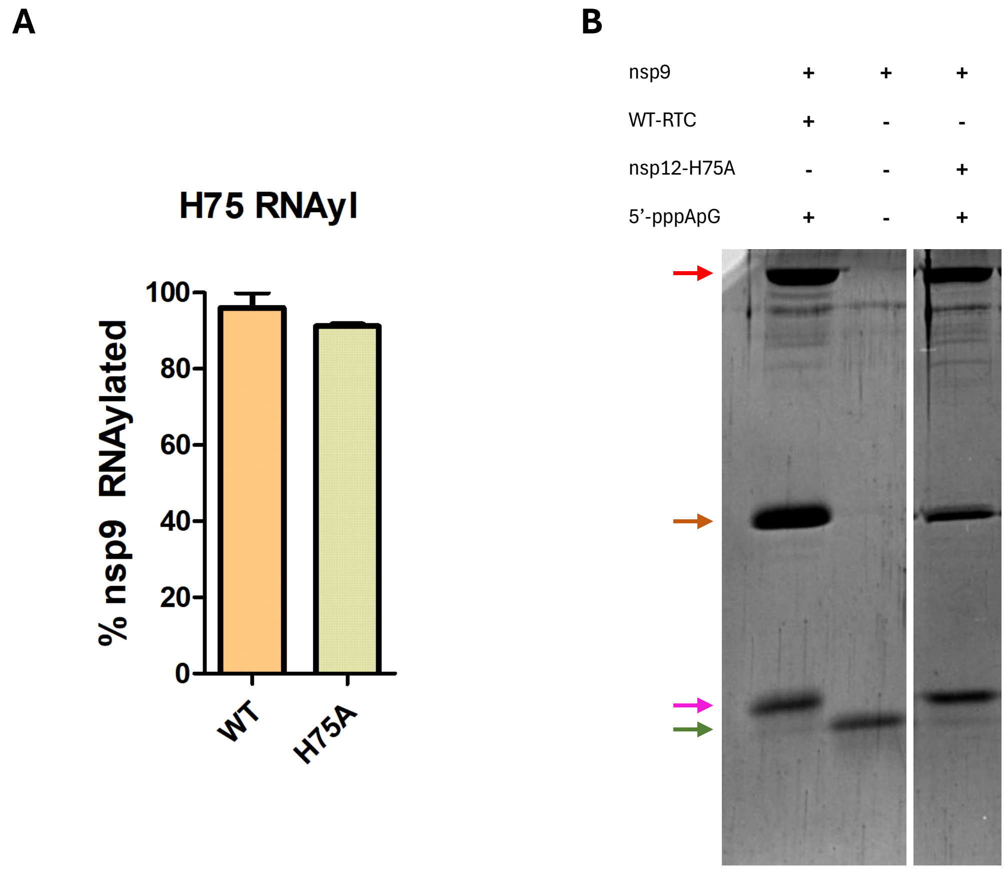


**Figure S10:** (A) RNAylation activity by WT-RTC and RTC containing nsp12-H75A mutant using 5'-pppApG. Reaction products were analyzed as in fig. 1F from three replicates. (B) Representative cropped SDS-PAGE gel image for nsp9 RNAylation experiment carried out with WT-RTC and RTC containing nsp12-H75A. Different protein bands of interest are marked using arrows – red for nsp12, brown for nsp8, green for nsp9 and pink for 5'-pApG-nsp9. All the images are representatives of three independent replicates.

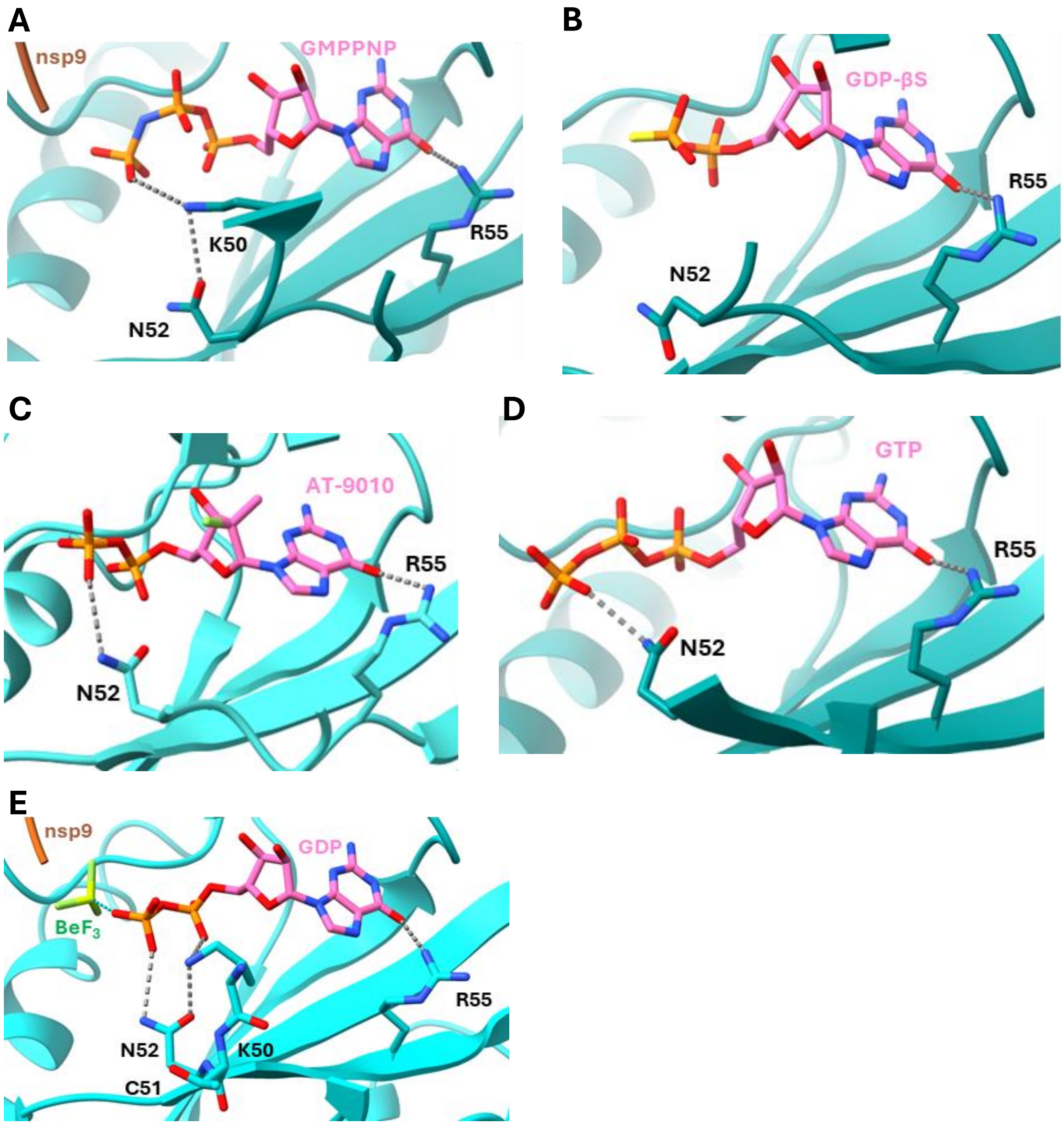


**Figure S11:** (A) Cartoon representation of interacting residues and GMPPNP in base-in pose in NiRAN active site (PDB ID 8GWE). (B) Cartoon representation of interacting residues and GDP-βS in base-in pose in NiRAN active site (PDB ID 8SQK). (C) Cartoon representation of interacting residues and AT-9010-DP (GDP analog) in base-in pose in NiRAN active site (PDB ID 7ED5). (D) Cartoon representation of interacting residues and GTP in base-in pose in NiRAN active site (PDB ID 7UOB). (E) Cartoon representation of interacting residues and GDP-BeF3 in base-in pose in NiRAN active site (PDB ID 9IKZ). For **A-E** interacting residues and nucleotides are shown as sticks and interactions are denoted by grey dashed lines

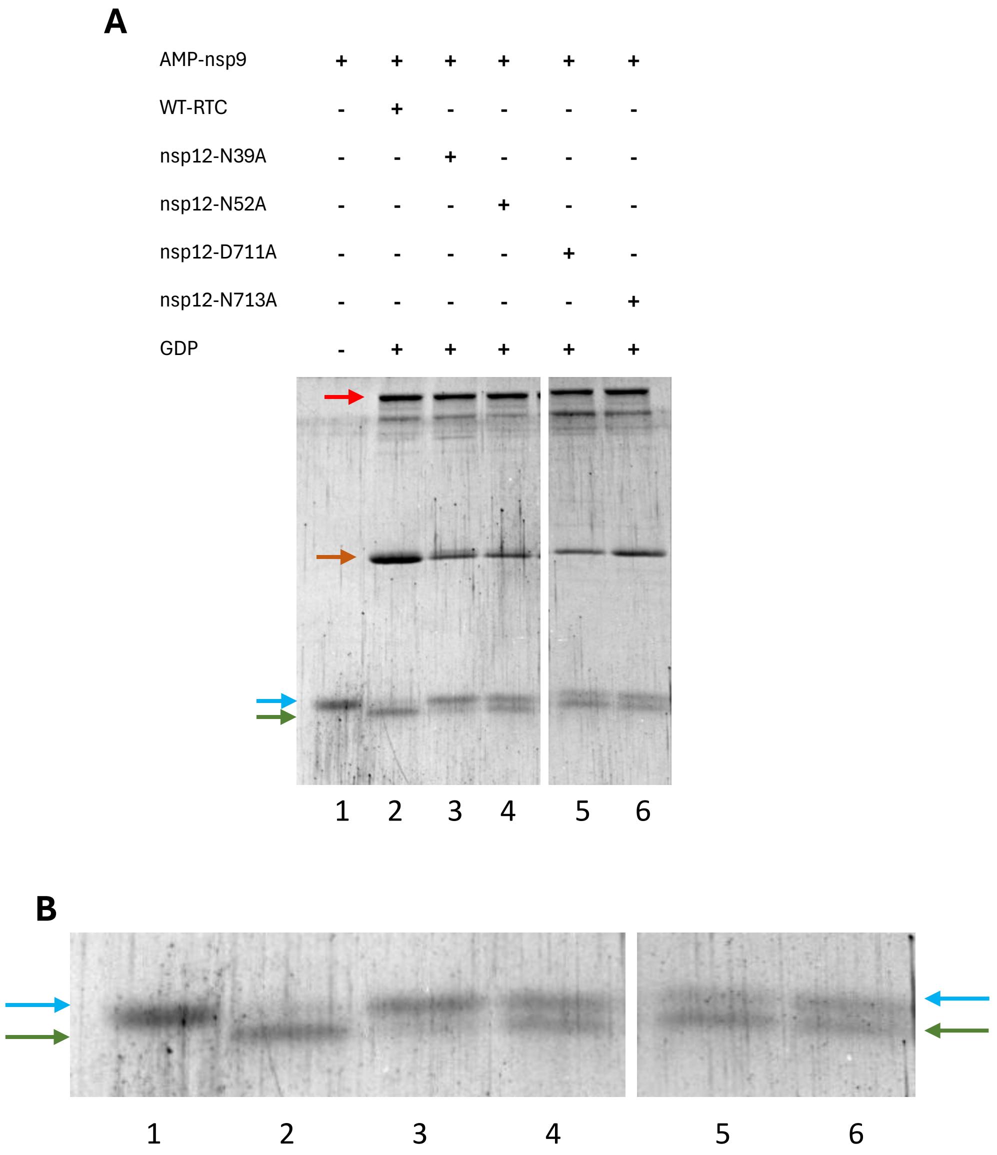


**Figure S12:** (A) Representative gel corresponding to fig. 3H. Cropped SDS-PAGE gel images for AMP-nsp9 deAMPylation experiment done with the WT-RTC and RTC containing indicated mutants of nsp12 using GDP shown along with no RNA control. Different protein bands of interest are marked using arrows – red for nsp12, brown for nsp8, green for nsp9 and blue for AMP-nsp9. Gels are cropped to show only lanes of interest. (B) Magnified and cropped image of the gels to highlight the modifications to nsp9. The numbers below the magnified images are lane numbers as in (A). Images are representative of atleast two independent replicates.

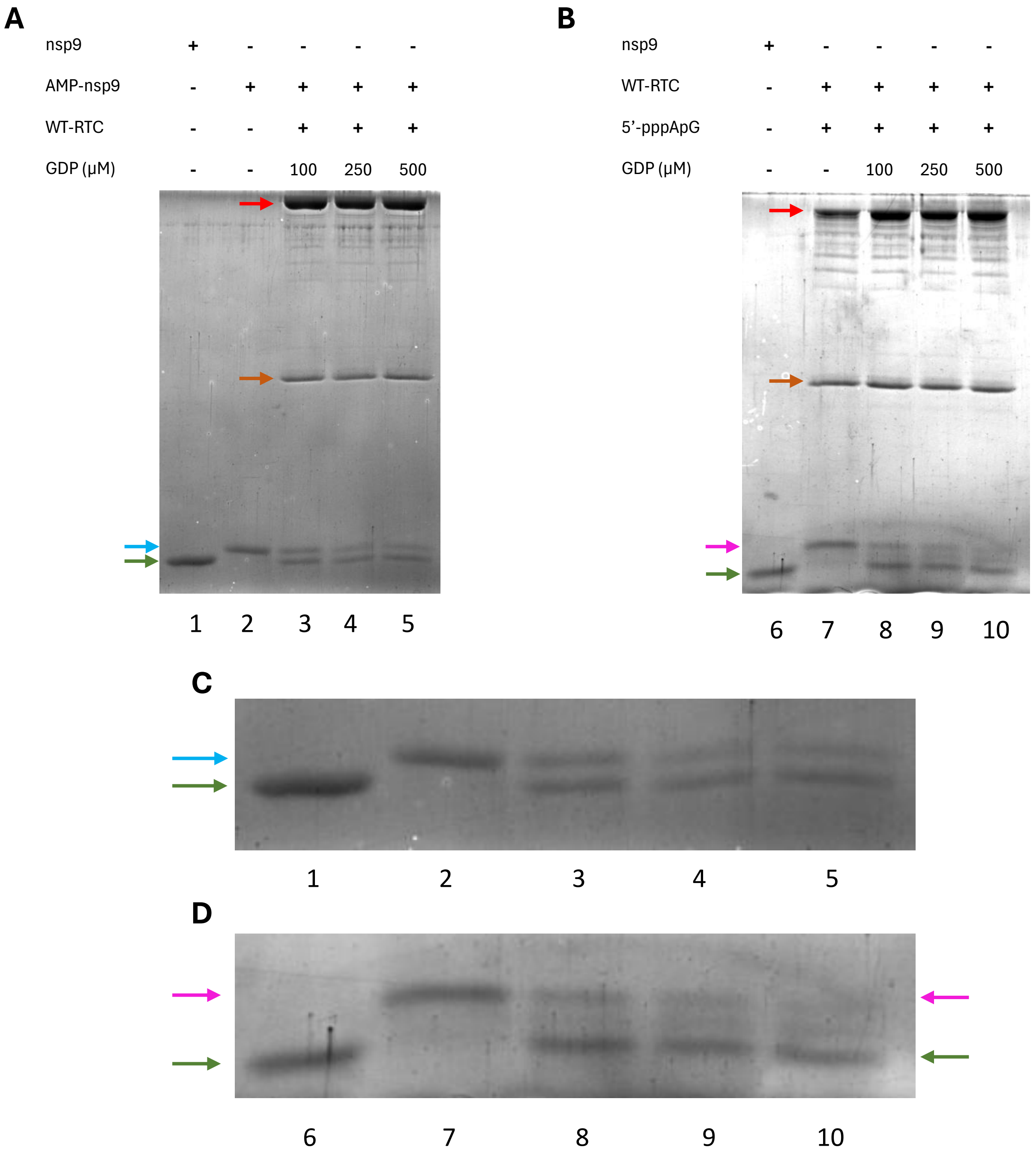


**Figure S13:** (A) Representative gel corresponding to fig. 4A. Cropped SDS-PAGE gel images for AMP-nsp9 deAMPylation experiment done with the WT-RTC using varying GDP concentrations as shown along with no RNA control. Different protein bands of interest are marked using arrows – red for nsp12, brown for nsp8, green for nsp9 and blue for AMP-nsp9. Gels are cropped to show only lanes of interest. Images are representative of three independent replicates. (B) Representative gel corresponding to fig. 4B. Cropped SDS-PAGE gel images for 5'-pApG-nsp9 deRNAylation experiment done with the WT-RTC using varying GDP concentrations as shown along with no RNA control. Different protein bands of interest are marked using arrows – red for nsp12, brown for nsp8, green for nsp9 and pink for 5'-pApG-nsp9. Gels are cropped to show only lanes of interest. Images are representative of three independent replicates. (C-D) Magnified and cropped image of the gels to highlight the modifications to nsp9. The numbers below the magnified images are lane numbers as in (A and B).

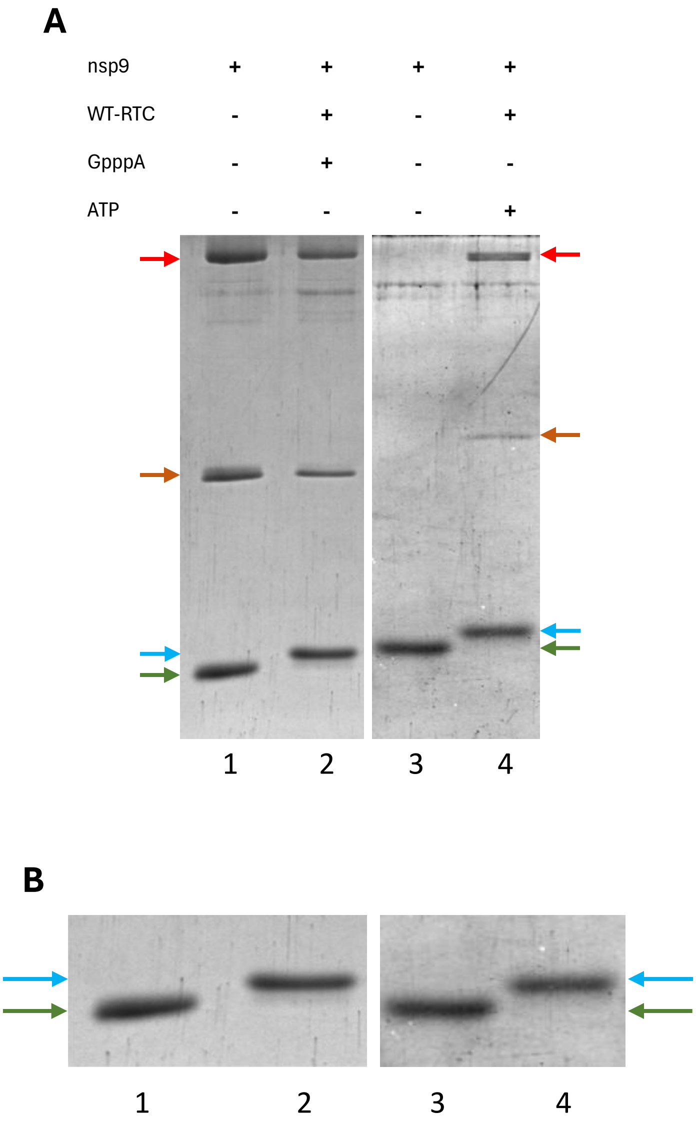


**Figure S14:** (A) Representative gel corresponding to fig. 4C. Cropped SDS-PAGE gel images for nsp9 NMPylation experiment done with the WT-RTC using ATP and GpppA shown along with no nucleotide control. Different protein bands of interest are marked using arrows – red for nsp12, brown for nsp8, green for nsp9 and blue for NMP-nsp9. Gels are cropped to show only lanes of interest. Images are representative of three independent replicates. (B) Magnified and cropped image of the gels to highlight the modifications to nsp9. The numbers below the magnified images are lane numbers as in (A).

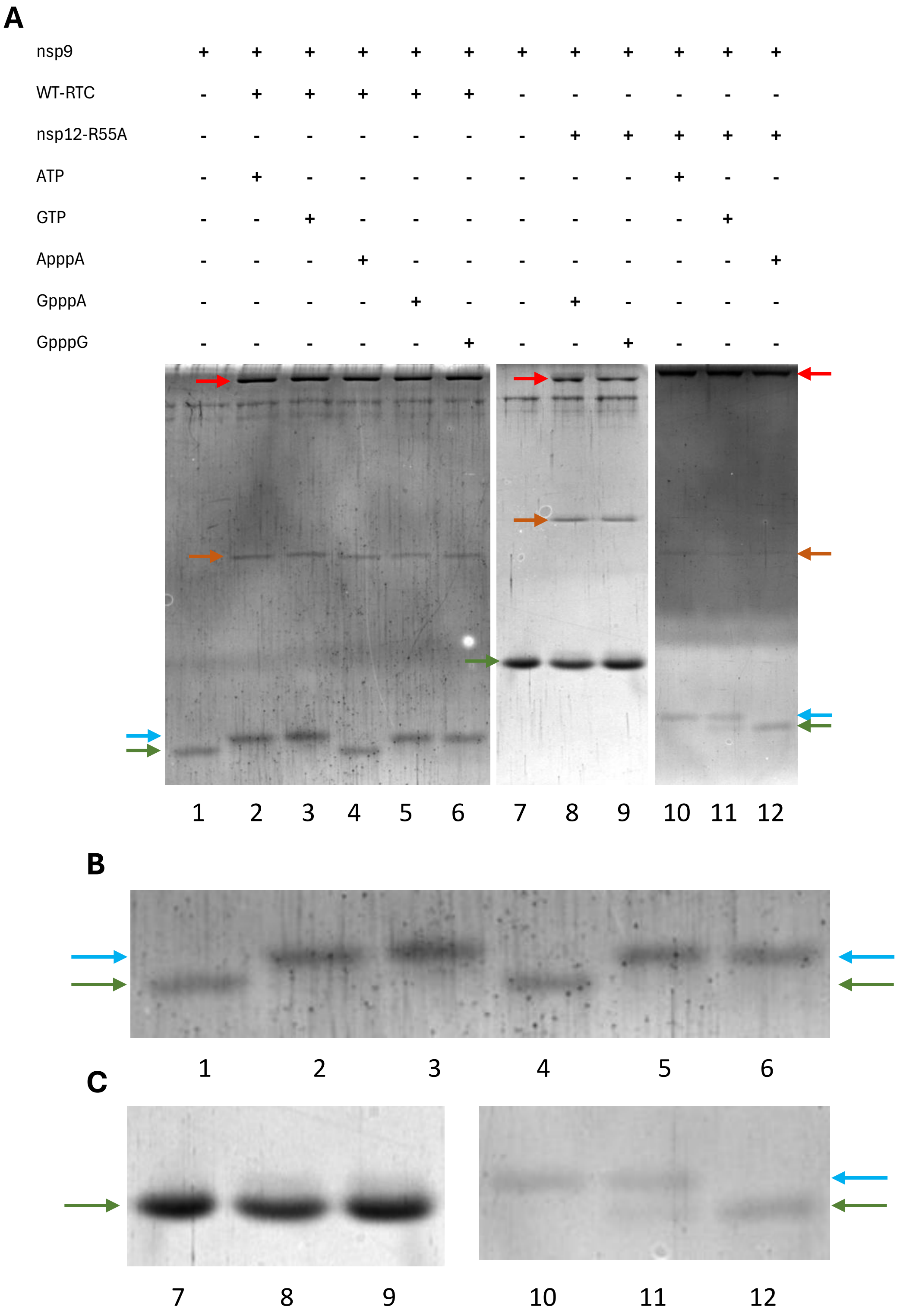


**Figure S15:** Representative gel corresponding to fig. 4D. Cropped SDS-PAGE gel images for nsp9 NMPylation experiment done with the WT-RTC and RTC containing indicated mutants of nsp12 using GpppA, GpppG and ApppA shown along with no nucleotide control. Different protein bands of interest are marked using arrows – red for nsp12, brown for nsp8, green for nsp9 and blue for NMP-nsp9. Gels are cropped to show only lanes of interest. (B-C) Magnified and cropped image of the gels to highlight the modifications to nsp9. The numbers below the magnified images are lane numbers as in (A). Images are representatives of three independent replicates.

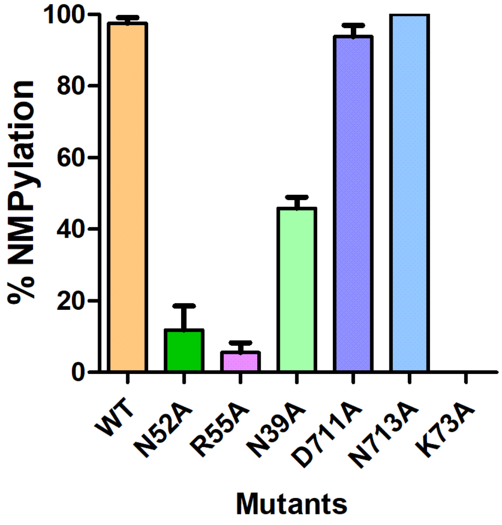


**Figure S16:** NMPylation activity by WT-RTC and RTC containing indicated mutants of nsp12 using GpppG. Reaction products were analyzed as in fig. 1D from three independent replicates.

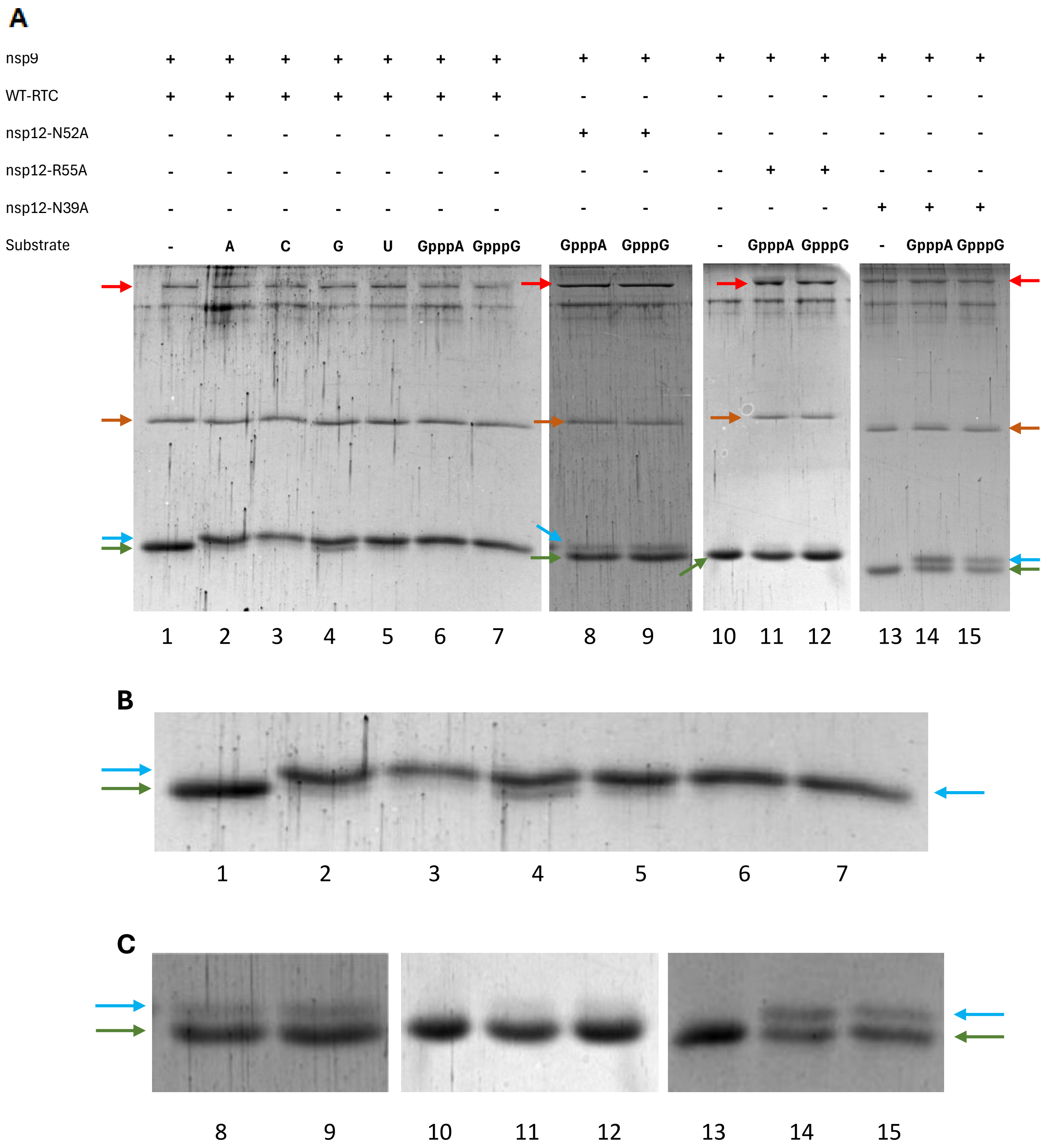


**Figure S17:** (A) Representative gel corresponding to fig. S14 and 4I. Cropped SDS-PAGE gel images showing nsp9 NMPylation using GpppA and GpppG with the WT-RTC and RTC containing indicated mutants of nsp12. Different protein bands of interest are marked using arrows – red for nsp12, brown for nsp8, green for nsp9 and blue for NMP-nsp9. Gels are cropped to show only lanes of interest. (B-C) Magnified and cropped image of the gels to highlight the modifications to nsp9. The numbers below the magnified images are lane numbers as in (A). Images are representatives of three independent replicates.

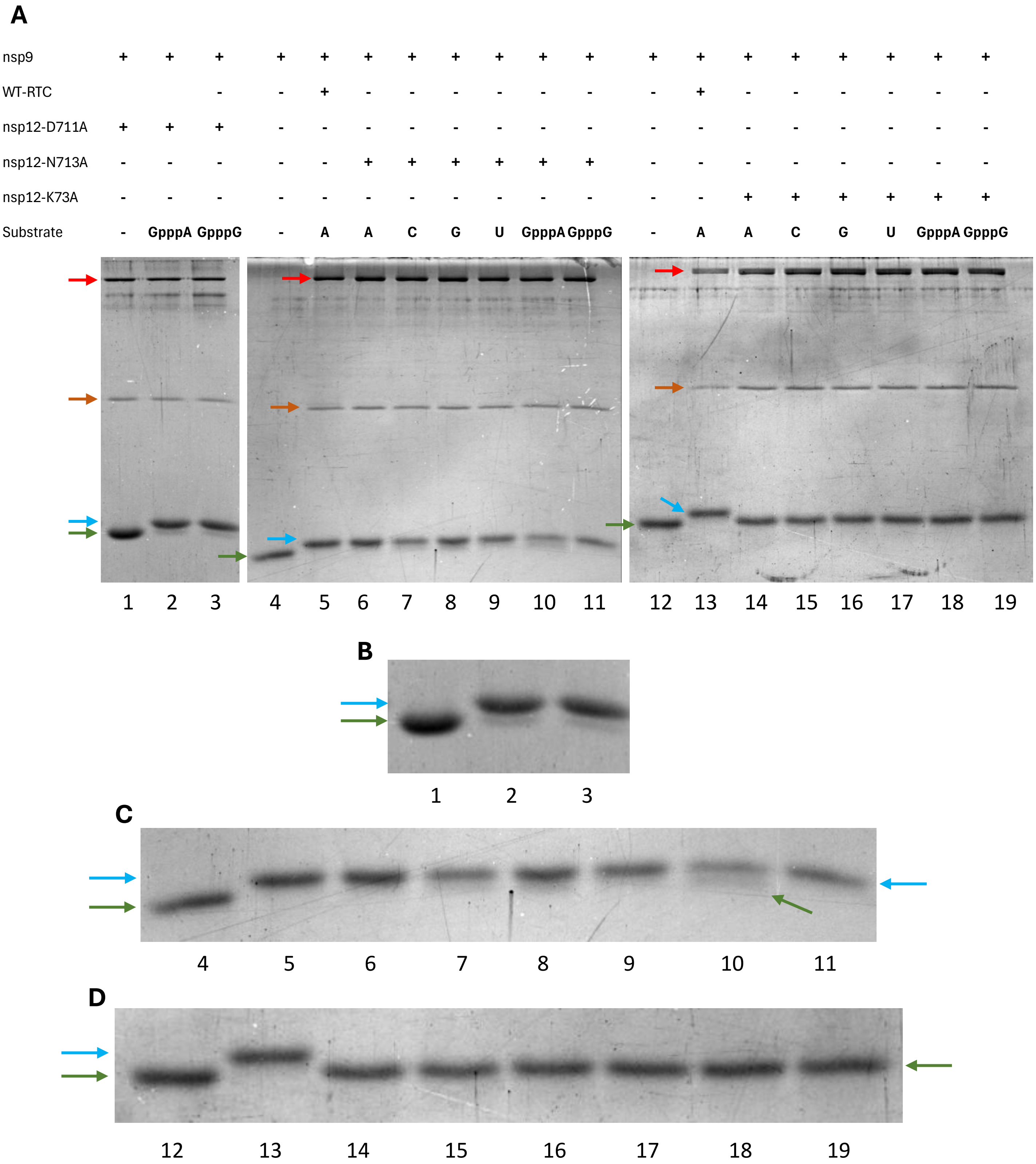


**Figure S18:** (A) Representative gel corresponding to fig. S14 and 4I. Cropped SDS-PAGE gel images showing nsp9 NMPylation using GpppA and GpppG with the WT-RTC and RTC containing indicated mutants of nsp12. Different protein bands of interest are marked using arrows – red for nsp12, brown for nsp8, green for nsp9 and blue for NMP-nsp9. Gels are cropped to show only lanes of interest. (B-D) Magnified and cropped image of the gels to highlight the modifications to nsp9. The numbers below the magnified images are lane numbers as in (A). Images are representatives of three independent replicates.

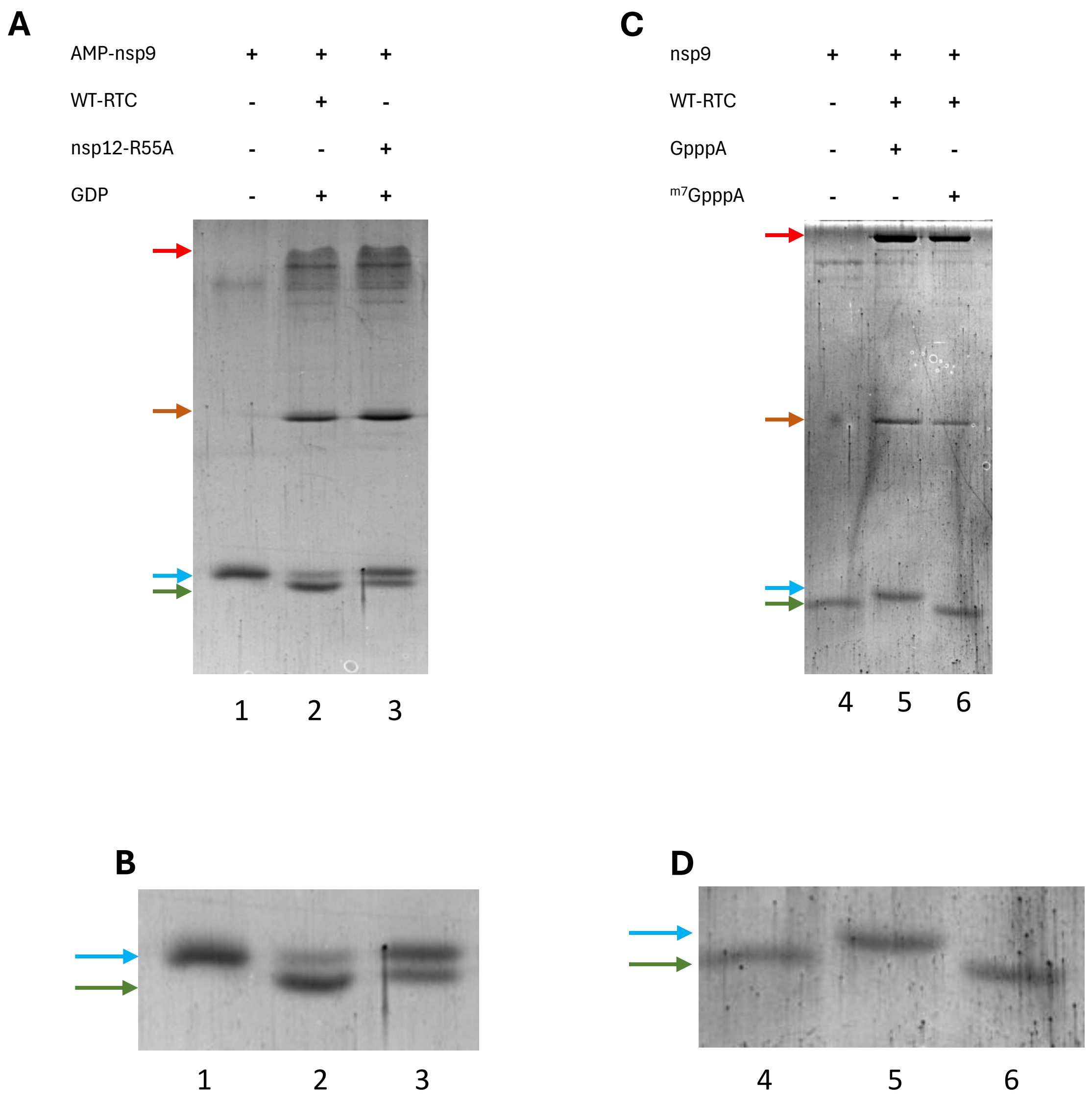


**Figure S19:** (A) Representative gel corresponding to fig. 4G. Cropped SDS-PAGE gel images for AMP-nsp9 deAMPylation experiment done with the WT-RTC and RTC containing indicated mutants of nsp12 using GDP shown along with no RNA control. Different protein bands of interest are marked using arrows – red for nsp12, brown for nsp8, green for nsp9 and blue for AMP-nsp9. Gels are cropped to show only lanes of interest. Images are representative of atleast two independent replicates. (B) Representative gel corresponding to fig. 4H. Cropped SDS-PAGE gel images for nsp9 NMPylation experiment done with the WT-RTC using GpppA and $^{m7}$GpppA shown along with no nucleotide control. Gels are cropped to show only lanes of interest. Images are representative of three independent replicates. (C-D) Magnified and cropped image of the gels to highlight the modifications to nsp9. The numbers below the magnified images are lane numbers as in (A). Images are representatives of at least two independent replicates.

| | | | | | | | | |
|---|---|---|---|---|---|---|---|---|
| nsp9 | + | + | + | + | + | + | + | + |
| WT-RTC | - | + | - | - | - | - | - | - |
| nsp12-N39A | - | - | + | - | - | - | - | - |
| nsp12-D711A | - | - | - | + | - | - | - | - |
| nsp12-N713A | - | - | - | - | + | - | - | - |
| nsp12-N52A | - | - | - | - | - | + | - | - |
| nsp12-R116A | - | - | - | - | - | - | + | - |
| nsp12-K73A | - | - | - | - | - | - | - | + |
| 5’-pppApG | - | - | - | - | - | - | - | - |

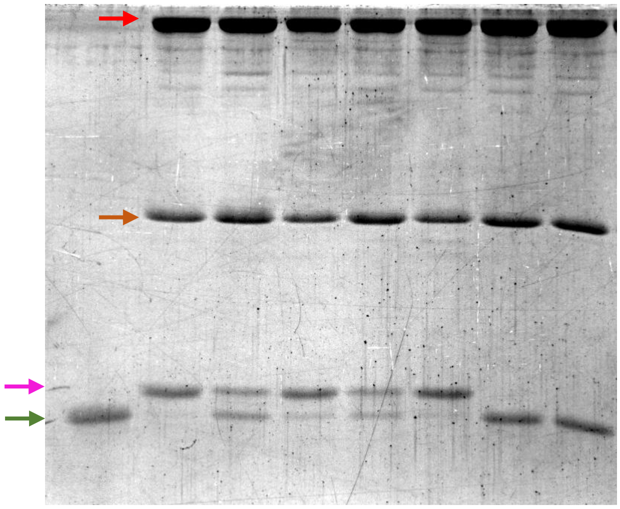

**Figure S20:** Representative cropped SDS-PAGE gel images for nsp9 RNAylation experiment done with WT-RTC and RTC containing indicated mutants of nsp12 using 5’-pppApG RNA shown along with no nucleotide control which shows nsp12-Lys73 has a catalytic role in RNAylation by NiRAN. Different protein bands of interest are marked using arrows – red for nsp12, brown for nsp8, green for nsp9 and blue for NMP-nsp9. Gels are cropped to show only lanes of interest. Images are representative of three independent replicates.

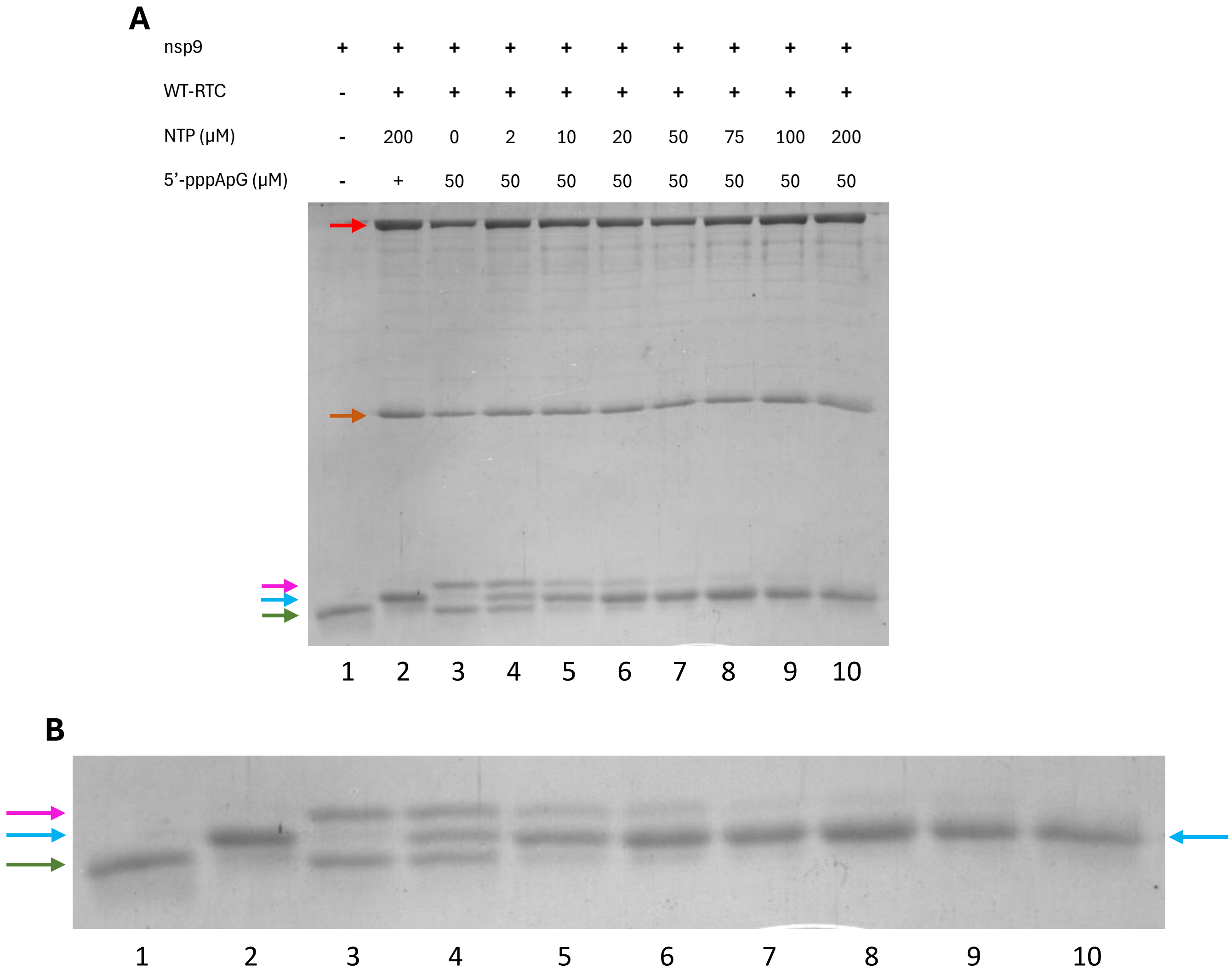


**Figure S21:** (A) Representative gel corresponding to fig. 5A. Cropped SDS-PAGE gel images for competition of nsp9 RNAylation using 5'-pppApG and NMPylation using varying concentration of NTP mix experiment done with the WT-RTC shown along with no nucleotide control. Different protein bands of interest are marked using arrows – red for nsp12, brown for nsp8, green for nsp9, blue for NMP-nsp9 and pink for5'-pApG-nsp9. Gels are cropped to show only lanes of interest. (B) Magnified and cropped image of the gels to highlight the modifications to nsp9. The numbers below the magnified images are lane numbers as in (A). Images are representative of three independent replicates.

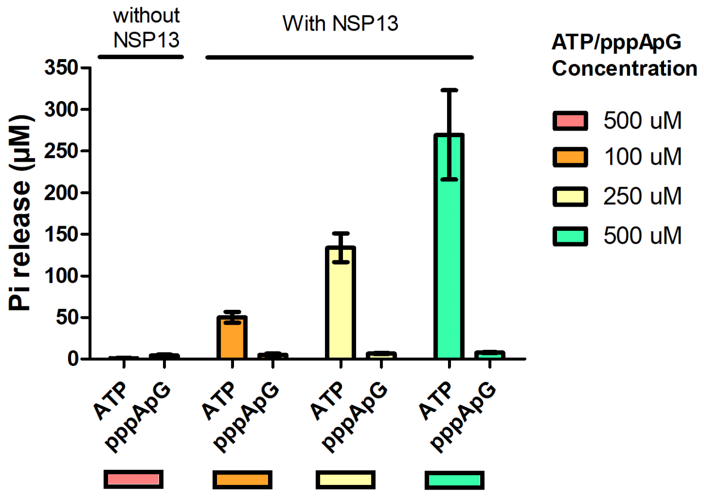


**Figure S22:** Nucleotide triphosphatase assay showing $P_i$ release from varying concentrations of ATP and pppApG using WT-nsp13. Malachite green assay was used to determine the $P_i$ release and the results were compared to an appropriate phosphate standard curve to obtain inorganic phosphate release. The error bars represent the standard deviation of the mean across three independent experiments.

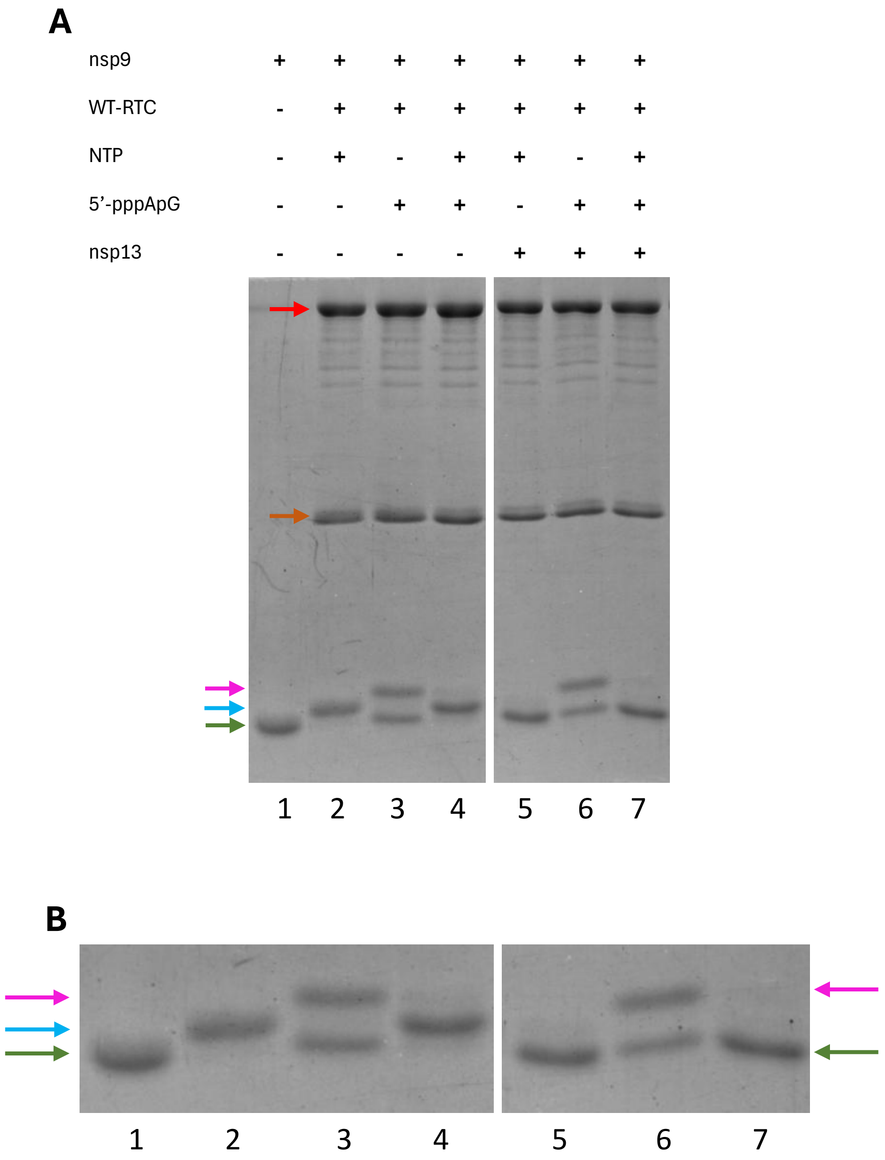


**Figure S23:** (A) Representative gel corresponding to fig. 5B. Cropped SDS-PAGE gel images for competition of nsp9 RNAylation using 5'-pppApG and NMPylation using NTP mix experiment done with the WT-RTC in the presence of nsp13 shown along with no nucleotide control. Different protein bands of interest are marked using arrows – red for nsp12, brown for nsp8, green for nsp9, blue for NMP-nsp9 and pink for5'-pApG-nsp9. Gels are cropped to show only lanes of interest. (B) Magnified and cropped image of the gels to highlight the modifications to nsp9. The numbers below the magnified images are lane numbers as in (A). Images are representative of one replicate.

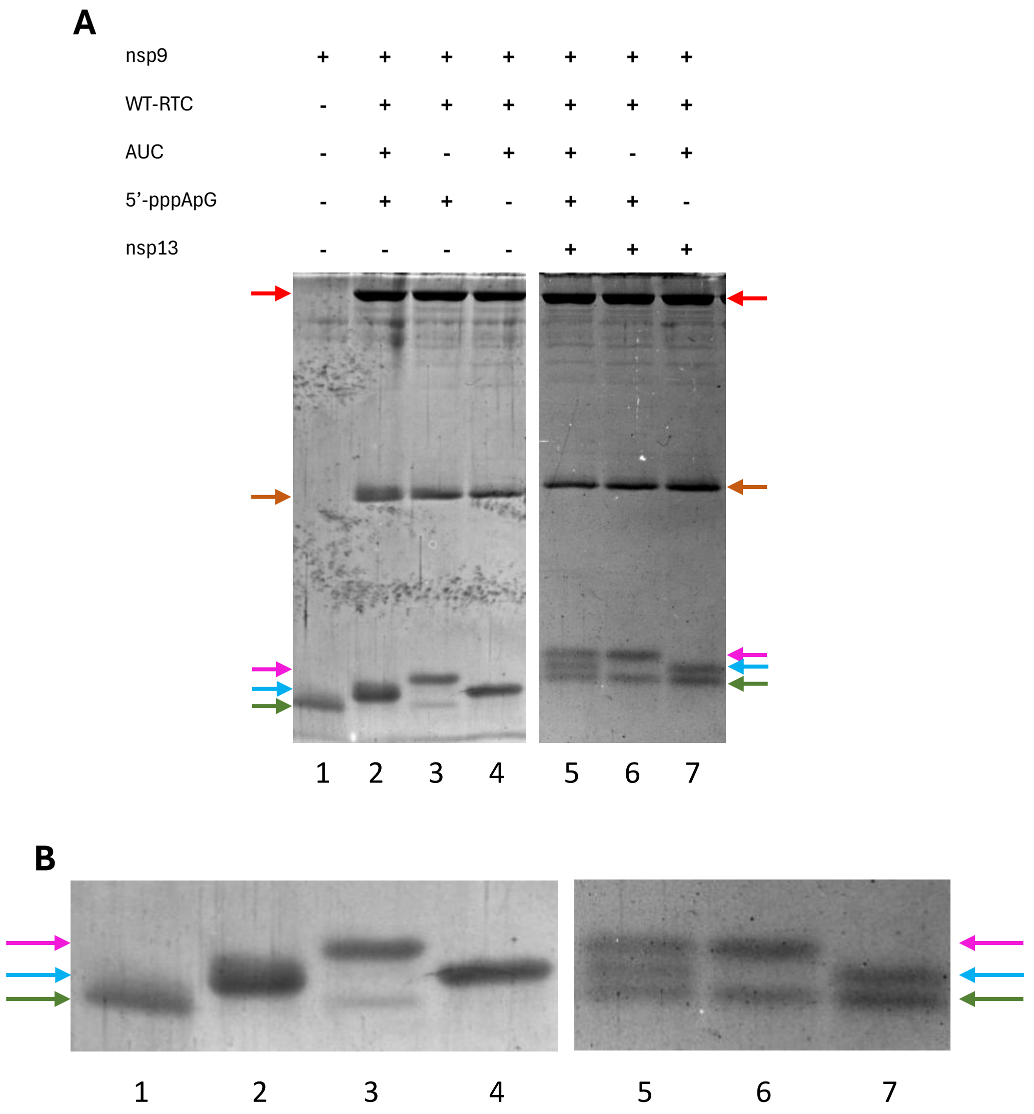


**Figure S24:** (A) Representative gel corresponding to fig. 5C. Cropped SDS-PAGE gel images for competition of nsp9 RNAylation using 5'-pppApG and NMPylation using AUC mix experiment done with the WT-RTC in the presence of nsp13 shown along with no nucleotide control. Different protein bands of interest are marked using arrows – red for nsp12, brown for nsp8, green for nsp9, blue for NMP-nsp9 and pink for5'-pApG-nsp9. Gels are cropped to show only lanes of interest. (B) Magnified and cropped image of the gels to highlight the modifications to nsp9. The numbers below the magnified images are lane numbers as in (A). Images are representative of three independent replicates.

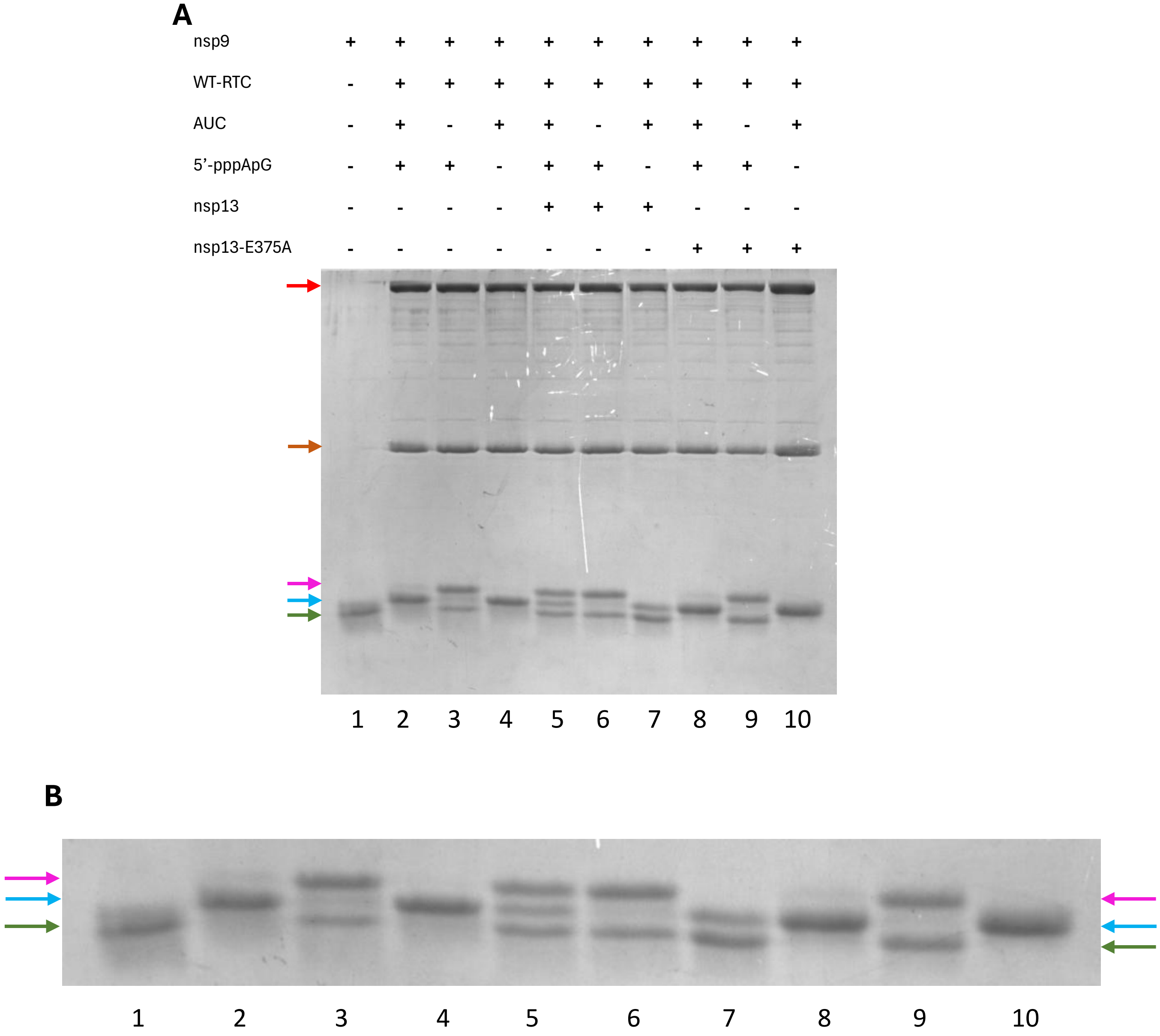


**Figure S25:** (A) Representative gel corresponding to fig. 5D. Cropped SDS-PAGE gel images for competition of nsp9 RNAylation using 5’-pppApG and NMPylation using AUC mix experiment done with the WT-RTC in the presence of nsp13 and nsp13-E375A shown along with no nucleotide control. Different protein bands of interest are marked using arrows – red for nsp12, brown for nsp8, green for nsp9, blue for NMP-nsp9 and pink for5’-pApG-nsp9. Gels are cropped to show only lanes of interest. (B) Magnified and cropped image of the gels to highlight the modifications to nsp9. The numbers below the magnified images are lane numbers as in (A). Images are representative of two independent replicates.

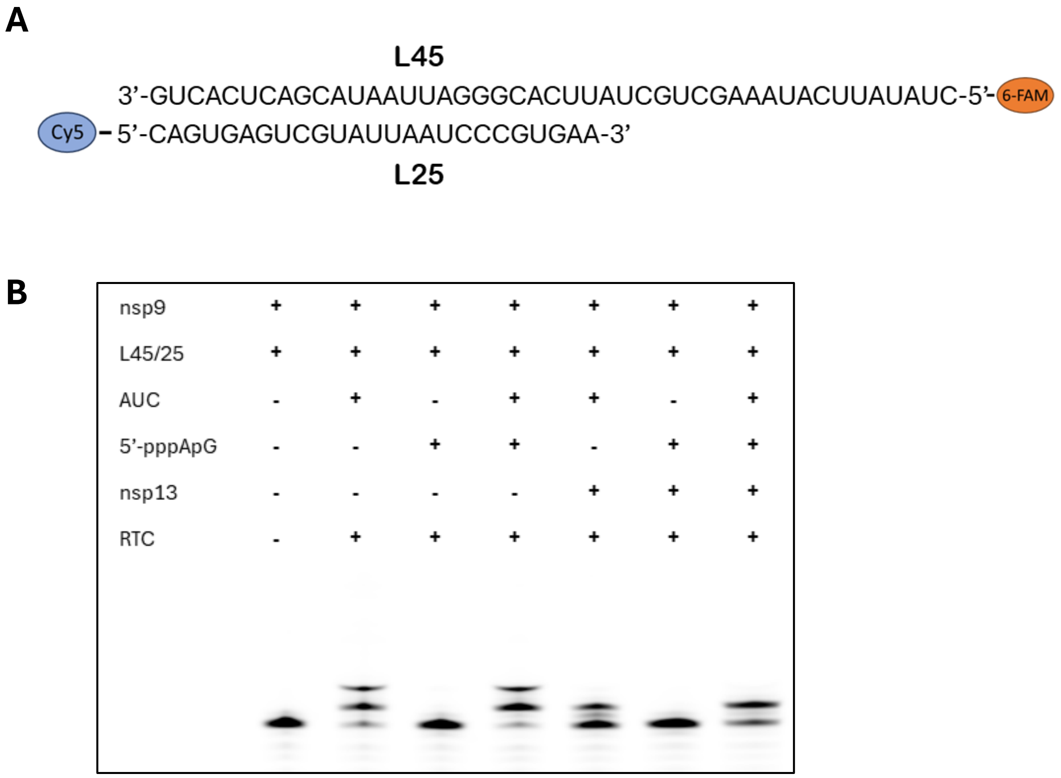


**Figure S26:** (A) Schematic showing L45/L25 RNA used for RTC polymerase activity. (B) Polymerase activity of WT-RTC using 200µM NTPs (All four - ATP, GTP, CTP and UTP) and 50µM pppApG, in the presence and absence of nsp13. The reaction products were analyzed using Urea-PAGE and the bands were detected using fluorescently labelled RNA. The image is representative of two independent replicates.

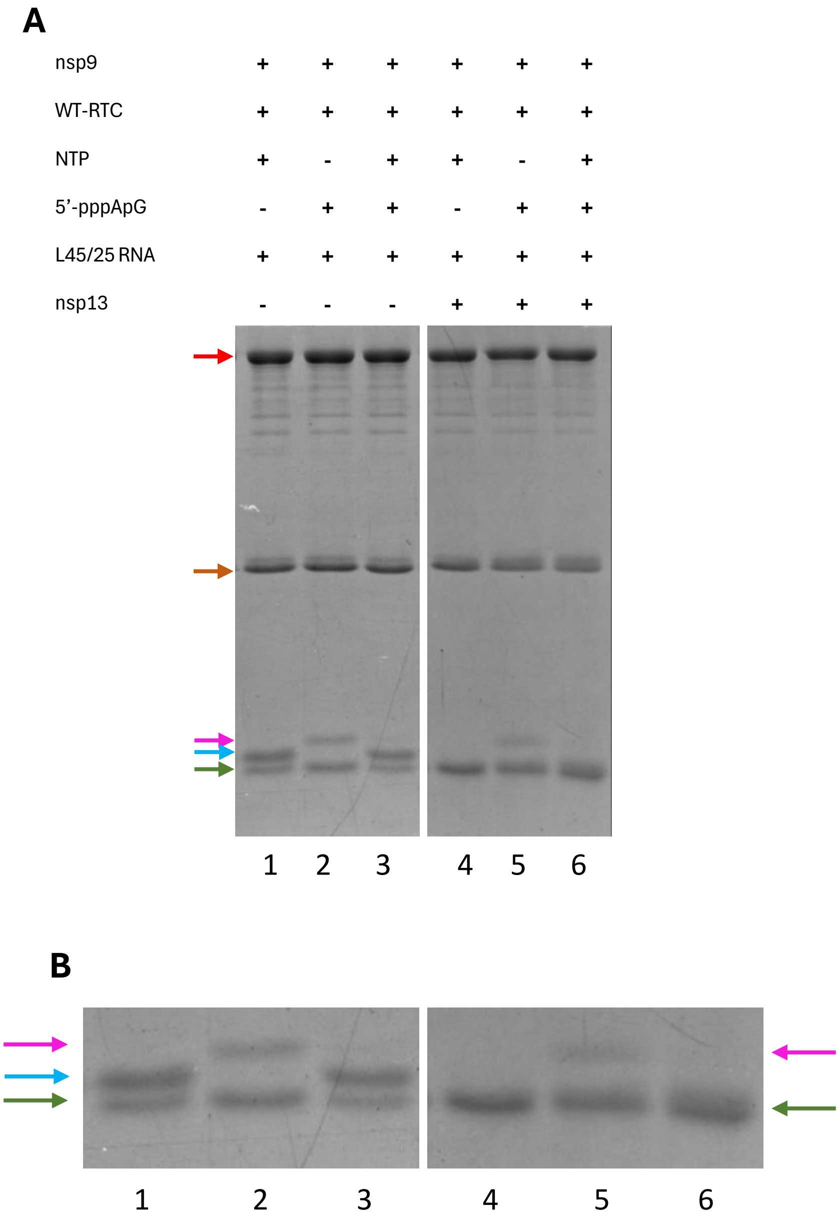


**Figure S27:** (A) Representative gel corresponding to fig. 5E. Cropped SDS-PAGE gel images for competition of nsp9 RNAylation using 5’-pppApG and NMPylation using NTP mix experiment done with the WT-RTC in the presence of L45/25 RNA and nsp13 shown along with no nucleotide control. Different protein bands of interest are marked using arrows – red for nsp12, brown for nsp8, green for nsp9, blue for NMP-nsp9 and pink for5’-pApG-nsp9. Gels are cropped to show only lanes of interest. (B) Magnified and cropped image of the gels to highlight the modifications to nsp9. The numbers below the magnified images are lane numbers as in (A). Images are representative of one replicate.

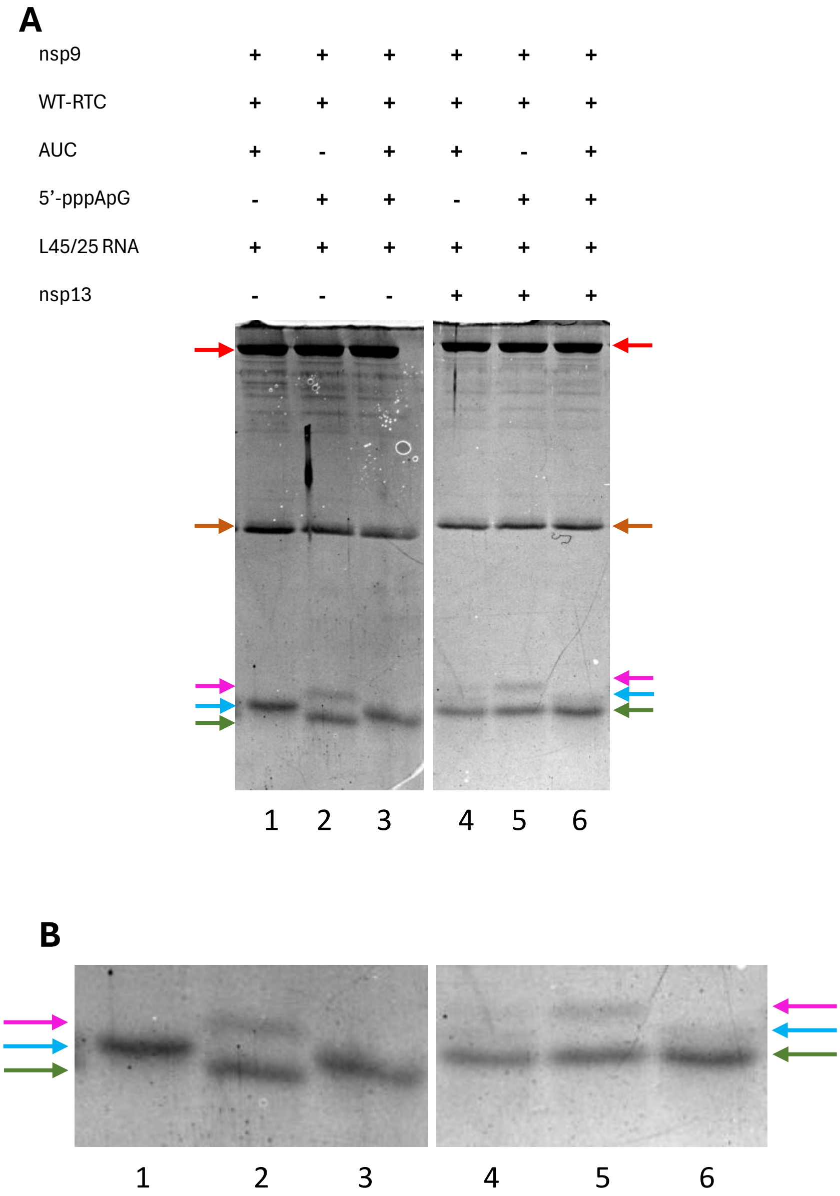


**Figure S28:** (A) Representative gel corresponding to fig. 5F. Cropped SDS-PAGE gel images for competition of nsp9 RNAylation using 5'-pppApG and NMPylation using AUC mix experiment done with the WT-RTC in the presence of L45/25 RNA and nsp13 shown along with no nucleotide control. Different protein bands of interest are marked using arrows – red for nsp12, brown for nsp8, green for nsp9, blue for NMP-nsp9 and pink for5'-pApG-nsp9. Gels are cropped to show only lanes of interest. (B) Magnified and cropped image of the gels to highlight the modifications to nsp9. The numbers below the magnified images are lane numbers as in (A). Images are representative of three independent replicates.

**Table S1: Table of PDB structures with nucleotide bound in the base-in pose in NiRAN**

| PDBID | Protein Components | NiRAN Ligand | Features | Reference |
|---|---|---|---|---|
| 8SQK | nsp7, $nsp8_2$, nsp9 & nsp12 | RNA-nsp9 + GDPβS | Asn52 oriented away from nucleotide; Lys50 interacts with β-phosphate | Small et al., 2023 |
| 7ED5 | nsp7, $nsp8_2$ & nsp12 | AT-9010-DP | Asn52 oriented towards nucleotide with no interaction; Lys50 interacts with β-phosphate | Shannon et al., 2022 |
| 7UOB | nsp7, $nsp8_2$ & nsp12 | GTP | Asn52 interacts with the phosphates of the base, not with Lys50 | Malone et al., 2023b |
| 8GWE | nsp7, $nsp8_2$, nsp9, nsp12 & $nsp13_2$ | RNA-nsp9 + GMPPNP | Asn52 does not interact with the phosphates of the base, but interacts with Lys50 | Yan et al., 2022 |
| 8GWF | nsp7, $nsp8_2$, nsp9, nsp12 & $nsp13_2$ | RNA-nsp9 (no density) + GTP | | Yan et al., 2022 |
| 8GWI | nsp7, $nsp8_2$, nsp9, nsp12 & $nsp13_2$ | SMP-nsp9 + GTP | | Yan et al., 2022 |
| 8GWN | nsp7, $nsp8_2$, nsp9, nsp12 & $nsp13_2$ | ATMP-nsp9 (no density) + GMPPNP | | Yan et al., 2022 |
| 8GWO | nsp7, $nsp8_2$, nsp9, nsp12 & $nsp13_2$ | UMP-nsp9 + GMPPNP | | Yan et al., 2022 |
| 8GWM | nsp7, $nsp8_2$, nsp9, nsp12 & $nsp13_2$ | MMP-nsp9 + GMPPNP | | Yan et al., 2022 |
| 8GWK | nsp7, $nsp8_2$, nsp9, nsp12 & $nsp13_2$ | RMP-nsp + GMPPNP | | Yan et al., 2022 |
| 9UHT | nsp7, $nsp8_2$, nsp9, nsp12 & $nsp13_2$ | RNA-nsp9 + GMPPNP | | Huang et al., 2025 |
| 9IKZ | nsp7, $nsp8_2$, nsp9, nsp12 & $nsp13_2$ | RNA-nsp9 + GDP-$BeF_3^-$ | Asn52 interacts with the γ-phosphates of the base and Lys50 | Huang et al., 2025 |
| 8GWG | nsp7, $nsp8_2$, nsp9, nsp12 & $nsp13_2$ | SMP-nsp9 + GMPPNP | Asn52 interacts with Lys50. Both do not face the nucleotide | Yan et al., 2022 |

**Table S2: Table of PDB structures with nucleotide bound in the base-out pose in NiRAN**

| PDBID | Protein Components | NiRAN Ligand | Features | Reference |
|---|---|---|---|---|
| 7KRO | nsp7, $nsp8_2$, nsp12 & $nsp13_2$ | ADP | Base in parallel stack interaction with His75 | B. Malone et al., 2021 |
| 7KRP | nsp7, nsp8 & nsp12 | ADP | | B. Malone et al., 2021 |
| 7KRN | nsp7, $nsp8_2$, nsp12 & nsp13 | ADP | | B. Malone et al., 2021 |
| 7RE0 | nsp7, $nsp8_2$, nsp12 & $nsp13_2$ | ADP | | Chen et al., 2022 |
| 7RE1 | nsp7, $nsp8_2$, nsp12 & $nsp13_2$ | ADP | | Chen et al., 2022 |
| 7RE2 | nsp7, $nsp8_2$, nsp12 & nsp13 | ADP | | Chen et al., 2022 |
| 7RE3 | (nsp7, $nsp8_2$, nsp12 & $nsp13_2)_2$ | ADP | | Chen et al., 2022 |
| 7RDX | nsp7, $nsp8_2$, nsp12 & $nsp13_2$ | ADP | | Chen et al., 2022 |
| 7RDY | nsp7, $nsp8_2$, nsp12 & $nsp13_2$ | ADP | | Chen et al., 2022 |
| 7RDZ | nsp7, $nsp8_2$, nsp12 & $nsp13_2$ | ADP | | Chen et al., 2022 |
| 6XEZ | nsp7, $nsp8_2$, nsp12 & $nsp13_2$ | ADP | | Chen et al., 2020 |
| 7CYQ | nsp7, $nsp8_2$, nsp9, nsp12 & $nsp13_2$ | GDP | Base in T-stack interaction with His75 | Yan et al., 2021 |
| 9IMM | nsp7, nsp82, nsp9, nsp12 & $nsp13_2$ | 5'-pppA-RNA | Base away from His75 | Yan et al., 2025 |
| 9IMK | (nsp7, nsp82, nsp9, nsp12 & $nsp132)_2$ | 5'-pppA-RNA | | Yan et al., 2025 |

**Table S3: Table of PDB structures with nucleotide bound in the base-up pose in NiRAN**

| PDBID | Protein Components | NiRAN Ligand | Features | Reference |
|---|---|---|---|---|
| 8SQK | nsp7, $nsp8_2$, nsp9 & nsp12 | RNA-nsp9 + GDPβS | 1st base interacts with Asp711 and Asn713 | Small et al., 2023 |
| 8SQJ | nsp7, $nsp8_2$, nsp9 & nsp12 | RNA-nsp9 | | Small et al., 2023 |
| 8GWB | nsp7, $nsp8_2$, nsp9, nsp12 & $nsp13_2$ | RNA-nsp9 | 1st base interacts with Asn39 and Asn713 | Yan et al., 2022 |
| 8GWE | nsp7, $nsp8_2$, nsp9, nsp12 & $nsp13_2$ | RNA-nsp9 + GMPPNP | | Yan et al., 2022 |
| 9UHT | nsp7, $nsp8_2$, nsp9, nsp12 & $nsp13_2$ | RNA-nsp9 + GMPPNP | | Huang et al., 2025 |
| 8SQ9 | nsp7, $nsp8_2$, nsp9 & nsp12 | UMPCPP | Pre-catalysis; Base interacts with Asn39 and Asn713 | Small et al., 2023 |
| 8GWG | nsp7, $nsp8_2$, nsp9, nsp12 & $nsp13_2$ | SMP-nsp9 + GMPPNP | Post-catalysis; Base interacts with Asn39 and Asn713 | Yan et al., 2022 |
| 8GWI | nsp7, $nsp8_2$, nsp9, nsp12 & $nsp13_2$ | SMP-nsp9 + GTP | | Yan et al., 2022 |
| 8GWO | nsp7, $nsp8_2$, nsp9, nsp12 & $nsp13_2$ | UMP-nsp9 + GMPPNP | | Yan et al., 2022 |
| 8GW1 | nsp7, $nsp8_2$, nsp9, nsp12 & $nsp13_2$ | UMP-nsp9 | | Yan et al. 2022 |
| 8GWM | nsp7, $nsp8_2$, nsp9, nsp12 & $nsp13_2$ | MMP-nsp9 + GMPPNP | | Yan et al., 2022 |
| 8GWK | nsp7, $nsp8_2$, nsp9, nsp12 & $nsp13_2$ | RMP-nsp9 + GMPPNP | | Yan et al., 2022 |
